# Inhibition of proteasomal degradation rescues a pathogenic variant of mitochondrial Respiratory chain assembly 1 factor

**DOI:** 10.1101/375725

**Authors:** Karthik Mohanraj, Michal Wasilewski, Cristiane Benincá, Dominik Cysewski, Jarosław Poznanski, Paulina Sakowska, Markus Deckers, Sven Dennerlein, Erika Fernandez-Vizarra, Peter Rehling, Michal Dadlez, Massimo Zeviani, Agnieszka Chacinska

## Abstract

Nuclear and mitochondrial genome mutations lead to various mitochondrial diseases, many of which affect the mitochondrial respiratory chain. The proteome of the intermembrane space (IMS) of mitochondria consists of several important assembly factors that participate in the biogenesis of mitochondrial respiratory chain complexes. The present study comprehensively analyzed a recently identified IMS protein, RESpiratory chain Assembly 1 (RESA1) factor, or cytochrome c oxidase assembly factor 7 (COA7) that is associated with a rare form of mitochondrial leukoencephalopathy and complex IV deficiency. We found that RESA1 requires the mitochondrial IMS import and assembly (MIA) pathway for efficient accumulation in the IMS. We also found that pathogenic mutant versions of RESA1 are imported slower than the wild type protein, and mislocalized mutant proteins are degraded in the cytosol by proteasome machinery. Interestingly, proteasome inhibition rescued both the mitochondrial localization of mutant RESA1 and complex IV activity in patient-derived fibroblasts. We propose that proteasome inhibition is a novel therapeutic approach for a broad range of mitochondrial pathologies that are associated with the excessive degradation of mitochondrial proteins that is caused by genetic mutations or biogenesis defects.

## Introduction

The constant import of nuclear-encoded mitochondrial proteins into mitochondria is required for healthy and functional mitochondria. Mitochondrial precursor proteins utilize various versatile import machineries for efficient import and proper localization into the destined sub compartments, including the outer membrane (OM), intermembrane space (IMS), inner membrane (IM), and matrix (Chacinska et al, 2009; Endo & Yamano, 2010; Neupert & Herrmann, 2007; Schmidt et al, 2010; Schulz et al, 2015; Wasilewski et al, 2017). Among the various import machineries, the mitochondrial intermembrane space import and assembly (MIA) pathway is responsible for the import and stable accumulation of cysteine-rich proteins in the IMS (Chacinska et al, 2004; Naoe et al, 2004; Terziyska et al, 2005). The identification and structural characterization of human MIA40, the central component of the MIA pathway, unveiled mechanistic details of this machinery in mammalian cells (Banci et al, 2009; Erdogan et al, 2018; Fischer et al, 2013; Hofmann et al, 2005; Sztolsztener et al, 2013). Similar to the yeast system, the human MIA pathway works in two steps. First, the reduced form of a precursor protein enters mitochondria *via* translocase of the outer membrane (TOM) and interacts with the hydrophobic cleft of MIA40 through the characteristic mitochondria IMS-sorting signal (MISS) / IMS-targeting signal (ITS) signal in its sequence (Lu et al, 2004; Milenkovic et al, 2009; Sideris et al, 2009). Secondly, this interaction is strengthened by intermolecular disulfide bond formation that ensures the oxidation of precursor proteins (Banci et al, 2010; Mesecke et al, 2005; Muller et al, 2008). The oxidation of precursor proteins leads to the reduction of cysteine residues in the CPC motif of MIA40. To facilitate the successive import cycle, the cysteine residues are oxidized by the transfer of electrons to the sulfhydryl oxidase human augmenter of liver regeneration (ALR). The chaperone-like trapping activity of Mia40 only, without disulfide bond formation, is also used to ensure the mitochondrial localization of some precursor proteins (Banci et al, 2010; Banci et al, 2009; Koch & Schmid, 2014; Peleh et al, 2016; Wrobel et al, 2013). The classic MIA40 substrates are mostly small proteins (< 20 kDa), such as TIMM8A and COX6B, with a specific arrangement of cysteine residues, such as CX_3_C or CX_9_C (Bourens et al, 2012; Koehler, 2004). MIA40 is also involved in the import of non-canonical substrates, which do not possess CX_3_C or CX_9_C signals in their sequence such as apurinic/apyrimidinic endonuclease (APE1), cellular tumor antigen p53, and mitochondrial calcium uptake 1 (MICU1) (Barchiesi et al, 2015; Petrungaro et al, 2015; Zhuang et al, 2013). Similarly, in the yeast *Saccharomyces cerevisiae*, Mia40 facilitates the import of non-canonical substrates, such as mitochondrial import inner membrane translocase subunit Tim22 and mitochondrial inner membrane protease Atp23 (Weckbecker et al, 2012; Wrobel et al, 2013). Thus, MIA40 is involved in the import of mitochondrial proteins with diverse topologies and architecture.

Mitochondria are multifunctional organelles that are involved in crucial biological processes, including respiration and apoptosis. Mitochondrial dysfunction is associated with many diseases (Costa & Scorrano, 2012; Gorman et al, 2016; Suomalainen & Battersby, 2018; Viscomi et al, 2015). Among these, the impaired biogenesis of MIA40 and its substrates contributes to a significant percentage of mitochondrial pathology (Friederich et al, 2017; Koehler et al, 1999; Roesch et al, 2002; Tranebjaerg et al, 2000). Interestingly, mutations of genes that encode MIA40 substrates (e.g., TIMM8A, COX6B and NDUFB10) are associated with the significant loss of proteins in patients, consequently leading to impairments of the biogenesis of many other IMS and IM proteins. Interestingly, many mitochondrial precursor proteins, before they productively reach the mitochondrial compartment, are under control of the ubiquitin-proteasome system (UPS), a major protein-degrading machinery that is involved in maintaining cellular protein homeostasis (Bragoszewski et al, 2013; Radke et al, 2008). Furthermore, IMS proteins can undergo reductive unfolding and slide back to the cytosol where they are also degraded by the UPS (Bragoszewski et al, 2015). However, the contribution of cytosolic protein control mechanisms to mitochondriopathies has not been investigated.

The present study identified a cysteine-rich IMS protein, RESA1/COA7 as a new non-canonical substrate of MIA40. RESA1 is a Sel1 repeat-containing protein that was previously shown to be involved in the assembly of respiratory chain complexes I and principally complex IV (Kozjak-Pavlovic et al, 2014; Martinez Lyons et al, 2016). We characterized the interaction between RESA1 and MIA40 and its import and localization in mitochondria. We also characterized pathogenic disease-causing RESA1 mutants (Martinez Lyons et al, 2016) as defective in their biogenesis and degraded in the cytosol by the UPS. Importantly, inhibition of the UPS system led to the partial rescue of defective RESA1 variants, thus suggesting a conceptually novel strategy for therapeutic interventions.

## Results

### RESA1: New MIA40-interacting protein

To identify new interacting partners/substrates of MIA40, we performed affinity purification of MIA40_FLAG_ that was overexpressed in human Flp-In T-REx293 cells and subjected the eluate fraction to mass spectrometry. Specificity was calculated as the ratio of the protein signal intensity in the bait purification (MIA40_FLAG_) to the protein signal intensity in the control purification. The identification of previously known MIA40-interacting proteins (ALR and AIF1) validated the specificity of our affinity purification. Among the proteins that co-purified with MIA40_FLAG_, we identified the cysteine-rich protein RESA1 with high abundance and specificity (Fig 1A) in three biological replicates. RESA1 is very distinct from classic MIA40 substrates due to the presence of 13 cysteine residues that are neither surrounded by classic MIA40-targeting signals (MISS/ITS) and nor arranged in classic motifs (CX_3_C or CX_9_C). Immunoblotting of the MIA40_FLAG_ eluate fraction with RESA1 antibody confirmed our mass spectrometry observation (Fig 1B, lane 4). As a positive control, we observed good co-purification of ALR and AIF in the eluate fraction (Fig 1B, lane 4). To verify the possible interaction between RESA1 and ALR (a known MIA40 partner), we performed affinity purification *via* ALR_FLAG_, and only a small amount of RESA1 co-purified with ALR (Fig 1C).

**Figure 1.**
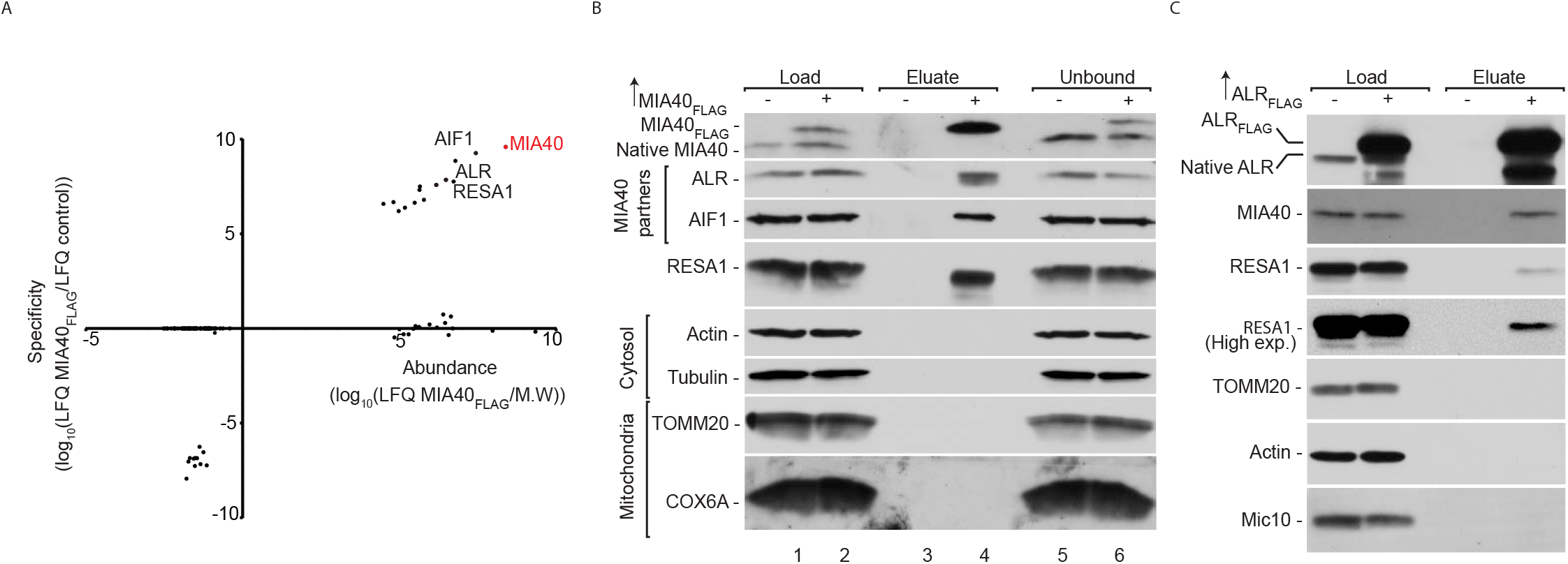
RESA1 interacts with MIA40. (A) Protein abundance is the normalized signal intensity (LFQ) for a protein divided by its molecular weight. Specificity (enrichment) is the ratio of the protein LFQ intensity in the MIA40_FLAG_ fraction to control samples. The LFQ for proteins that were not detected in the control samples was arbitrarily set to 1 for calculation purposes. LFQ, Label-Free-Quantification; M.W., molecular weight. (B) Flp-In 293 T-REx cells that expressed MIA40_FLAG_ were solubilized, and the affinity purification of MIA40_FLAG_ was performed. The fractions were analyzed by SDS-PAGE and Western blot. Load: 2.5%. Eluate: 100%. Unbound: 2.5%. (C) Flp-In 293 T-REx cells that expressed ALR_FLAG_ were solubilized, and the affinity purification of MIA40_FLAG_ was performed. The fractions were analyzed by SDS-PAGE and Western blot. Load: 2.5%. Eluate: 100%.

### RESA1 interacts with MIA40 through disulfide bonds

Since MIA40 interacts with many precursor proteins *via* disulfide bonds, we investigated whether the interaction with RESA1 involves the formation of disulfide bridges. We first performed affinity purification *via* MIA40_FLAG_ and probed the eluate fraction using MIA40 and RESA1 antibodies under reducing (dithiotheritol [DTT]) and non-reducing (iodoacetamide [IAA]) conditions. Immunoblotting against both RESA1 and MIA40 revealed a band of approximately 45 kDa (Fig 2A, lanes 2 and 3) under non-reducing conditions, which would match the combined molecular weight of a covalent, disulfide-bonded complex of MIA40_FLAG_ (15.9 kDa) and RESA1 (25.7 kDa). Therefore, we assumed that RESA1 may interact with MIA40 through disulfide bonds. We tested this hypothesis using mitochondria that were isolated from cells that expressed wild type and cysteine residue mutant versions of MIA40_FLAG_, namely C53S, C55S, and C53S-C55S (denoted SPS), under non-reducing conditions (Fig 2B). We observed two bands that represented native MIA40-RESA1 and MIA40_FLAG_-RESA1 complexes in cells that expressed wild type and C53S MIA40_FLAG_ (Fig 2B, lanes 2, 3, 7, and 8) and only one band that corresponded to the native MIA40-RESA1 complex in cells that expressed C55S MIA40_FLAG_ and SPS MIA40_FLAG_, respectively (Fig 2B, lanes 4, 5, 9, and 10). We also confirmed this observation in cellular protein extract that was probed with anti-RESA1 antibody. This indicated the existence of a covalently bound complex between MIA40 and RESA1 under *in vivo* conditions (Fig 2C). The second cysteine residue in the CPC motif of MIA40 was previously shown to be crucial for the interaction with classic MIA40 substrates (Banci et al, 2009). Therefore, we checked the cysteine dependency of RESA1 binding by performing affinity purification *via* an MIA40 cysteine mutant protein (C53S, C55S, and SPS) that was expressed in Flp-In T-REx 293 cells. We observed RESA1 only in the eluate fractions of wtMIA40_FLAG_ and C53S MIA40_FLAG_ (Fig 2D, lanes 6 and 7) and not in the other mutants (Fig 2D, lanes 9 and 10). Thus, the second cysteine of the CPC motif (i.e., C55) was crucial for the disulfide bonding of MIA40 with RESA1.

**Figure 2.**
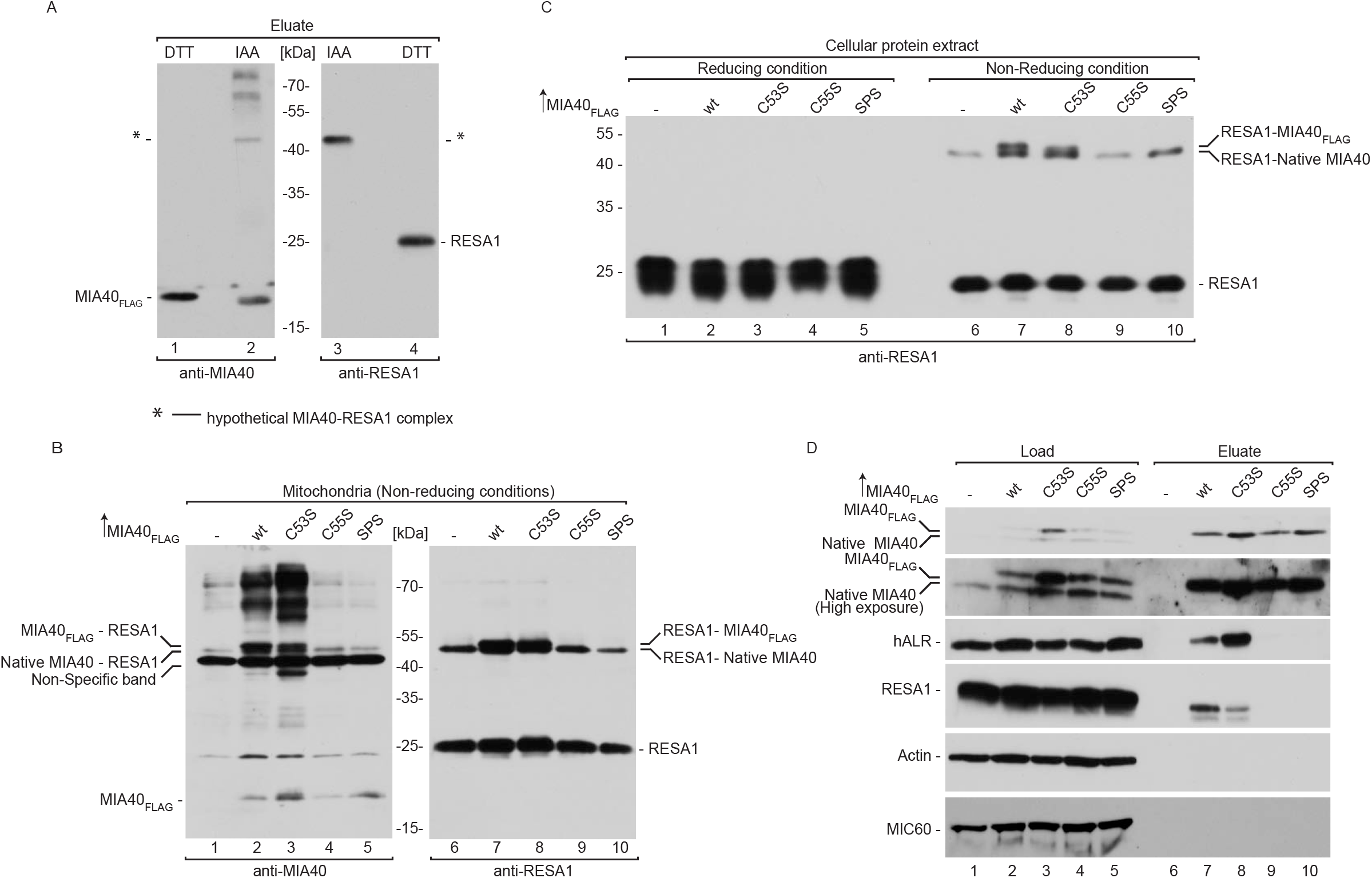
RESA1 interacts with MIA40 by disulfide bonding. (A) Flp-In 293 T-REx cells that expressed MIA40_FLAG_ were solubilized, and the affinity purification of MIA40_FLAG_ was performed. Eluate fractions were solubilized under reducing (DTT) or non-reducing (IAA) conditions and analyzed by SDS-PAGE and Western blot. DTT, dithiothreitol; IAA, iodoacetamide. Eluate: 100%. (B) Mitochondria that were isolated from cells that expressed MIA40_FLAG_ (wildtype and mutants as indicated) were solubilized under non-reducing conditions and analyzed by SDS-PAGE and Western blot. (C) Cellular protein extracts were isolated from Flp-In 293 T-REx cells that expressed wildtype and mutant forms of MIA40_FLAG_ under non-reducing conditions. The extract was analyzed by non-reducing SDS-PAGE and Western blot. (D) Cellular extracts from Flp-In 293 T-REx cells that expressed wildtype or mutant MIA40_FLAG_ were subjected to affinity purification. Load and eluate fractions were analyzed by reducing SDS-PAGE and Western blot. Load: 2.5%. Eluate: 100%.

### RESA1 exists as an oxidized protein in the IMS

RESA1 contains 13 cysteine residues. We evaluated the phylogenic diversity of cysteine residues in RESA1 orthologs in eukaryotes and found the strong conservation of 10 residues (C28, C37, C62, C71, C110, C111, C142, C150, C182, and C190) across eukaryotes, whereas three other cysteine residues (C24, C95, and C172) were present only in vertebrates (Fig EV1A). We hypothesized that the conserved cysteine residues may be involved in intramolecular disulfide bonds. To predict the cysteine residues that are involved in the disulfide bonds, we modeled the structure of RESA1 based on three proteins with high sequence similarity: Helicobacter cysteine rich protein C (1OUV), *Helicobacter pylori* cysteine rich protein B (4BWR), and a protective antigen that is present in *Escherichia coli* (1KLX). The modeled structure of RESA1 possessed five disulfide bonds with a unique pattern (disulfide bridges between cysteine residues that were separated by nine or 11 amino acids). The structure resembled those of the template proteins with subsequent helix-turn-helix (HTH) motifs that formed a super-helical structure that was analogous to previously reported structures. The N-terminal part of the protein did not have a proper template. Therefore, we modeled it *de novo* and manually adjusted the improperly modeled C24-C37 disulfide bridge to C28-C37. This modification was based on the following: (i) it fit the evolutionary conservation of cysteine residues, and (*ii*) formation of the C28-C37 bond concurs with the pattern of other disulfide bonds (i.e., cysteine separated by nine or 11 amino acids) in the protein. Hence, we proposed that the remaining three cysteine residues (C24, C95, and C172) were expected to be in a reduced state (Fig EV1B).

To validate our prediction of the RESA1 redox state, we employed direct and indirect thiol trapping assays on mitochondria that were isolated from human embryonic kidney 293 (HEK293) cells. The direct thiol trapping assay is based on the differential use of alkylating agents that bind free thiol residues in proteins and modify their migration in sodium dodecyl sulfate-polyacrylamide gel electrophoresis (SDS-PAGE). Binding of the low-molecular-mass agent IAA is neutral, whereas modification with 4-acetamido-4-maleimidylstilbene-2,2-disulfonic acid (AMS) alters the molecular mass of the protein by 0.5 kDa per each thiol residue. In the direct thiol assay, AMS modified the migration of RESA1 compared with IAA by approximately 1.5 kDa, which corresponds to three free thiol residues (Fig 3A, lanes 2 and 3). Therefore, we presumed that 10 cysteine residues would be involved in the disulfide bond, and the remaining three could be in a reduced state. To test this hypothesis, we performed an indirect thiol trapping assay that involved two-step thiol modification. Free thiols were first blocked with IAA, and the remaining cysteine residues that formed disulfide bridges were reduced with DTT and modified with AMS. The observed shift corresponded to cysteine residues that formed disulfide bridges in native RESA1 (Fig 3B, lane 4). As expected, we observed migration that corresponded to approximately 10 cysteine residues. As a control, we treated mitochondria directly with DTT to completely reduce all native disulfide bonds and then modified them with AMS (Fig 3B, lane 5). We observed higher migration of RESA1 compared with lane 4. These observations suggest that among the 13 cysteine residues, 10 are involved in disulfide bonds, and the other three exist in a reduced state. Based on the thiol trapping experiments and *in silico* modeling, we propose that RESA1 has five Sel-1 domain like repeats that are stabilized by disulfide bridges. The Sel-1 domains are characterized by a specific arrangement of cysteine residues (i.e., the fourth amino acid and fifth amino acid from the last amino acid in each domain is always a cysteine).

**Figure 3.**
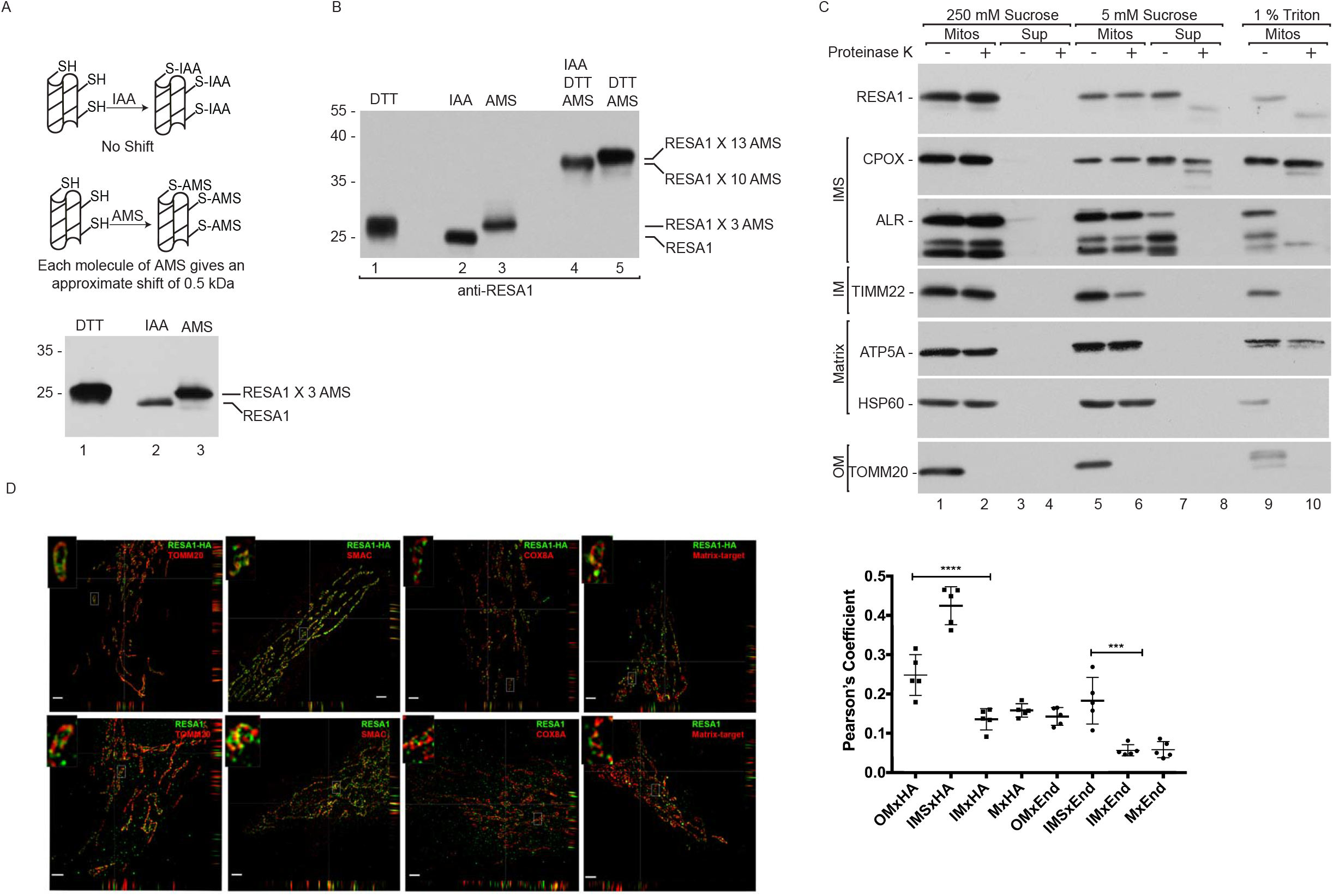
RESA1 exists as an oxidized protein in the intermembrane space of human mitochondria. (A) Schematic representation of the thiol trapping assay. Mitochondria were solubilized in sample buffer with either dithiothreitol (DTT), iodoacetamide (IAA), or 4-acetamido-4-maleimidylstilbene-2,2-disulfonic acid (AMS). The samples were analyzed by SDS-PAGE and Western blot. (B) Indirect thiol trapping assay. Mitochondria were pretreated with IAA as indicated to block free cysteine residues, and disulfide bonds were subsequently reduced by DTT. Finally, mitochondria were solubilized in sample buffer with AMS. (C) Localization of mitochondrial proteins analyzed by limited degradation by proteinase K in intact mitochondria (250 mM sucrose), mitoplasts (5 mM sucrose), and mitochondrial lysates (1% Triton X-100). The samples were analyzed by SDS-PAGE and Western blot. Mitos, mitochondria; Sup, post-mitochondria supernatant; OM, outer membrane; IM, inner membrane; IMS, intermembrane space. (D) The figure shows N-SIM super-resolution micrographs of one Z-stack (0.15 μm) orthogonal section (XYZ) of Hela cells or HeLa cells that stably expressed RESA1-HA transfected with different subcompartment markers TOMM20-DsRed (OM), COX8A-DsRed (IM), and mtPAGFP (Matrix-target) labeled with anti-HA, anti-RESA1 and Smac/Diablo (IMS) antibodies. The picture represents the majority population of cells from three independent experiments. Scale bar = 2 μm. The panel shows Pearson’s coefficient in a colocalized volume of different subcompartment combinations with anti-HA or anti-RESA1. The data are expressed as mean ± SD (n = 5). **p < 0.01, ***p < 0.001, ****p < 0.0001 (oneway ANOVA). M, matrix; End, endogenous RESA1; HA, RESA1-HA.

The mitochondrial IMS is the predicted destination for cysteine-rich oxidized protein that interacts with MIA40. To analyze the sub-mitochondrial localization of RESA1, we performed hypo-osmotic swelling of mitochondria that were isolated from HEK293 cells. In intact mitochondria, RESA1 was preserved from degradation, similar to other proteins that were localized inside mitochondria, such as ALR and coproporphyrinogen oxidase (CPOX) (IMS), TIMM22, and COX6B (IM), and HSP60 and ATP5A (matrix). As expected, the OM protein TOMM20 was efficiently degraded by proteinase K in intact mitochondria (Fig 3C, lanes 1-4). Upon rupturing the OM by swelling in hypotonic buffer, proteinase K degrades proteins that are exposed to the IMS, whereas matrix proteins remain protected. Accordingly, we observed the slight degradation of IM proteins (TIMM22) that faced the IMS side (Fig 3C, lanes 5-8). The matrix protein Hsp60 was completely resistant to proteinase K degradation, even upon swelling, and was degraded only by 1% Triton treatment (Fig 3C, lanes 9 and 10). Under these conditions, RESA1 was present in the supernatant fraction similarly to the soluble IMS proteins CPOX and ALR and was sensitive to proteinase K (Fig 3C, lanes 5-8). RESA1 has been previously reported to localize to the IMS or matrix (Kozjak-Pavlovic et al, 2014; Martinez Lyons et al, 2016). To further investigate the sub-mitochondrial localization of RESA1, we performed microscopy using the structured illumination technique (N-SIM). Several combinations of antibodies and targeted fluorescent proteins were used in order to verify the specificity of the observed co-localization. In particular we targeted fluorophores to a single mitochondrial sub-compartment and observed positive co-localization in the case of OM × OM (TOMM20 and TOMM20), IM × IM (SDHAand COX8A), and matrix × matrix (ACO2 and matrix target green fluorescent protein [GFP]) (Fig EV2). Conversely, we observed no co-localization when different subcompartments were stained: OM × IMS (TOMM20 and SMAC), OM × IM (SDHA and TOMM20), IM × matrix (COX8A and matrix target GFP), IMS × matrix (SDHA and MDH2), and OM × matrix (TOMM20 and matrix target GFP) (Fig EV2). These observations indicated that the technique provided sufficient resolution (~110 nm) to pair the correct subcompartment combinations of the positive controls TOMM20 × TOMM20 (OM × OM), COX8A × SDHA (IM × IM), and matrix-target GFP × ACO2 (matrix × matrix) from all of the combinations that were used as negative controls (Fig EV2). We then compared the signal of RESA1 with the markers of different mitochondrial subcompartments. We observed an increase in co-localization only with the IMS marker SMAC and not with markers of other subcompartments (Fig 3D). The IMS localization of RESA1 was similar with both endogenous RESA1 and HA-tagged overexpressed RESA1. Altogether, these findings indicate that RESA1 is a soluble protein of the IMS (Fig 3D).

### RESA1 does not influence the MIA pathway

The relatively strong interaction between RESA1 and MIA40 (see Fig 1 and 2) prompted us to explore the potential role of RESA1 in the MIA pathway. We first examined the steady-state levels of mitochondrial proteins upon RESA1 overexpression and found no changes in the levels of MIA40 or its dependent proteins, including ALR, COX6B, and TIMM22 (Fig 4A). We next investigated the influence of RESA1 on the import of MIA40 substrates by importing two substrates, TIMM8A (CX_9_C) and COX19 (CX_3_C), into mitochondria that has overexpressed levels of RESA1. We did not observe any significant difference in the import of TIMM8A or COX19 compared with the control (Fig 4B). These observations suggest that RESA1 overexpression does not influence either the steady-state levels of MIA40 or the import of MIA40 substrates.

**Figure 4.**
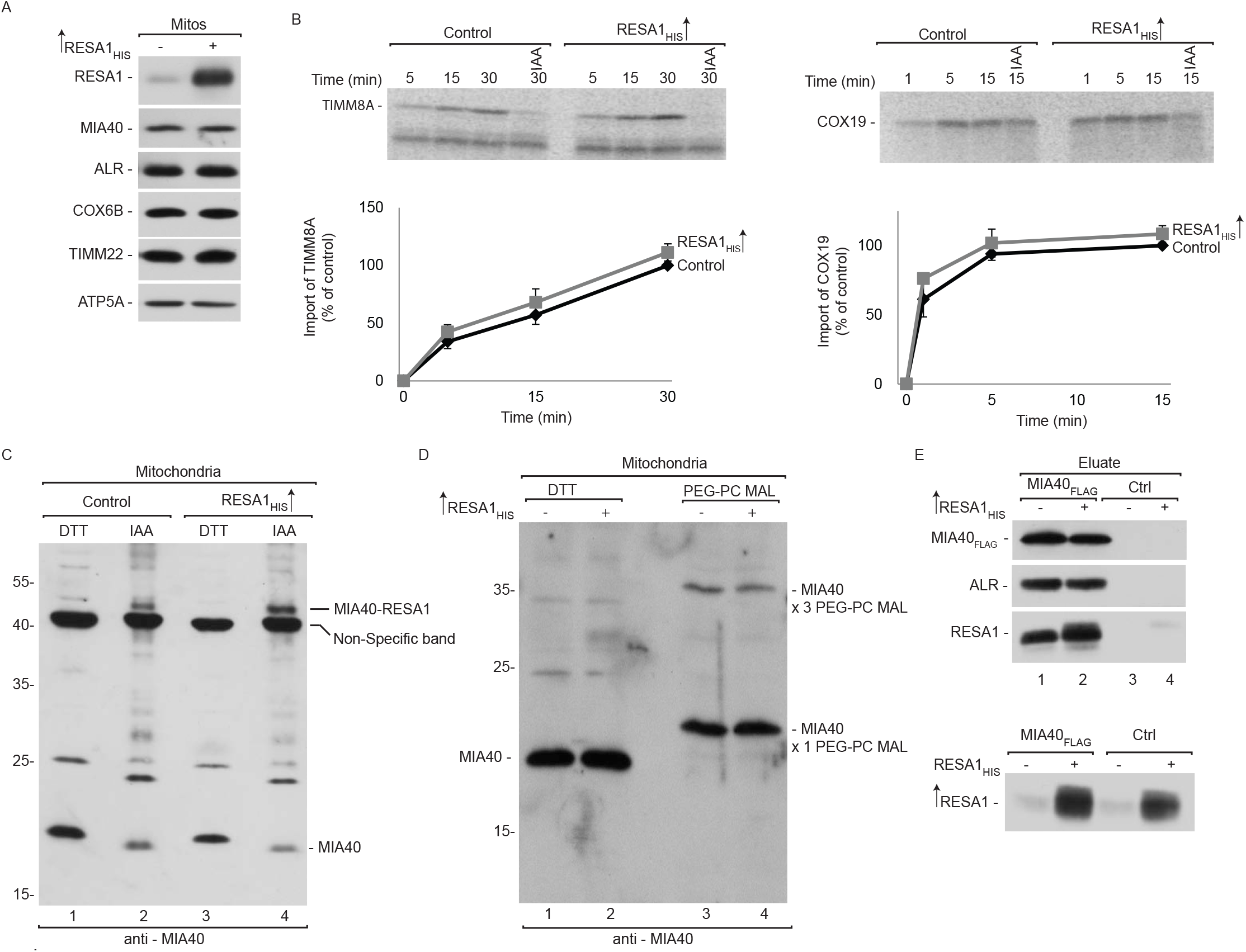
RESA1 does not influence the MIA pathway. (A) Mitochondria were isolated from cells that were transfected with a plasmid that encoded RESA1_HIS_ or an empty vector. Mitochondria were solubilized and analyzed by reducing SDS-PAGE and Western blot. Mitos, mitochondria. (B) Radiolabeled [^35^S]TIMM8A and [^35^S]COX19 precursors were imported into mitochondria that were isolated from cells that were transfected with a plasmid that encoded RESA1_HIS_ or an empty vector. The samples were analyzed by reducing SDS-PAGE and autoradiography. The results of three biological replicates were analyzed, quantified, and normalized to control mitochondria at 30 min. The data are expressed as mean ± SEM. IAA, iodoacetamide. (C) Mitochondria were isolated from cells that were transfected with a plasmid that encoded RESA1_HIS_ or an empty vector under reducing (DTT) and non-reducing (IAA) conditions and analyzed for levels of MIA40 by Western blot. (D) Mitochondria that were isolated from HEK293 cells were treated with either DTT or PEG-PC MAL. The samples were analyzed by SDS-PAGE and Western blot. (E) Flp-In 293 T-REx cells that expressed MIA40_FLAG_ were transfected with a plasmid that encoded RESA1_HIS_ or an empty vector. The affinity purification of MIA40_FLAG_ was performed, and eluate fractions were analyzed by SDS-PAGE and Western blot.

Cysteine residues in the CPC motif of MIA40 are crucial for the recognition and import of MIA40 substrates. We previously established that the second cysteine in the CPC motif of MIA40 is crucial for the interaction with RESA1 (Fig 2B-D). Therefore, we evaluated the levels of free MIA40 that is available for the import of substrates under nonreducing conditions. As expected, RESA1 overexpression did not significantly affect the level of free MIA40 (Fig 4C, lanes 2 and 4). We next checked the oxidation state of the cysteine residues in the CPC motif of MIA40 under conditions of RESA1 overexpression using the thiol-binding reagent PEG-PC MAL, which induces a shift of approximately 5 kDa per cysteine residue. The specificity of the observed bands was confirmed in cells that had genetically lower levels of MIA40 (Fig EV3A). We verified oxidation of the CPC motif of MIA40 in mitochondria that were isolated from cells that overexpressed RESA1. The treatment yielded two bands that were specific to MIA40. The lower band corresponded to the migration of MIA40 with PEG-PC MAL that was bound to one cysteine residue (C4). The second band corresponded to the binding of PEG-PC MAL to three cysteine residues (C4, C53, and C55). The CPC motif of MIA40 was oxidized to a similar extent in RESA1-overexpressed mitochondria and in the control mitochondria (Fig 4D, lanes 3 and 4). Thus, we confirmed that RESA1 overexpression did not affect the redox state of cysteine residues in the CPC motif, suggesting that both cysteine residues were equally available for importing MIA40 substrates, such as TIMM8A and COX6B. This observation complemented our previous finding that RESA1 overexpression did not affect the import of MIA40 substrates.

The role of ALR in the MIA pathway is well established, and the interaction between MIA40 and ALR is crucial for maintaining MIA40 activity (Banci et al, 2012; Sztolsztener et al, 2013). To investigate whether RESA1 influences this crucial interaction, we performed affinity purification *via* MIA40_FLAG_ to evaluate the MIA40-ALR interaction upon RESA1 overexpression. The levels of ALR that co-purified with MIA40 were similar in cells that overexpressed RESA1 and control cells (Fig 4E, lanes 1 and 2). We did not observe any difference in the MIA40-ALR interaction under RESA1-knockdown conditions (Fig EV3B). Consequently, RESA1 knockdown using two different oligonucleotides did not alter the levels of MIA40 or its substrates, such as COX6B, TIMM9, and TIMM22 (Fig EV3C). We also examined the MIA40-ALR interaction by affinity purification *via* ALR_FLAG_ under similar conditions. The amount of MIA40 that co-purified with ALR was similar under conditions of both RESA1 knockdown and overexpression (Fig EV3D, E). Thus, we conclude that RESA1 influences neither MIA40 pathway activity nor the import of MIA40 substrates to the IMS.

### MIA40 facilitates the import of RESA1 into mitochondria

To investigate whether MIA40 is involved in the import of RESA1, we performed *in organello* import of radiolabeled [^35^S] RESA1 precursor into mitochondria that were isolated from cells that overexpressed MIA40_FLAG_. We observed a significant increase in RESA1 import in mitochondria with MIA40_FLAG_ compared with the control (Fig 5A). As a positive control, import of the classic MIA40 substrate TIMM8A was significantly augmented by MIA40_FLAG_ overexpression (Fig 5B). Predictably, we also observed a significant increase in steady-state levels of RESA1, similar to other classic MIA40-dependent proteins (e.g., COX6B and ALR; Fig 5C). Thus, we conclude that RESA1 is imported by the MIA pathway and behaves like an MIA40 substrate. Considering the fact that ALR is an integral part of the MIA pathway, we also investigated whether ALR overexpression stimulates the import of RESA1. Contrary to MIA40 overexpression, the import of RESA1 was similar in ALR overexpressed mitochondria compared with control mitochondria (Fig 5D). Likewise, ALR overexpression did not alter the import of TIMM8A (Fig 5E). In agreement with the import results, steady-state levels of RESA1 and MIA40-dependent proteins did not increase upon ALR overexpression (Fig 5F). We also examined the influence of MIA40 depletion on RESA1 biogenesis. MIA40 knockdown using two different oligonucleotides decreased nearly 90% of MIA40 in HeLa cells (Fig EV4A). As expected, the levels of RESA1 were significantly reduced, similar to ALR and other substrates (e.g., COX6B and TIMM9). Proteins that are unrelated to the MIA pathway, such as actin, GAPDH, and ATP5A, were unchanged in MIA40 knockdown cells. This further substantiated the role of MIA40 in the biogenesis of RESA1. Interestingly, ALR knockdown also affected the steady state levels of RESA1 and MIA40-dependent proteins (Fig EV4B).

**Figure 5.**
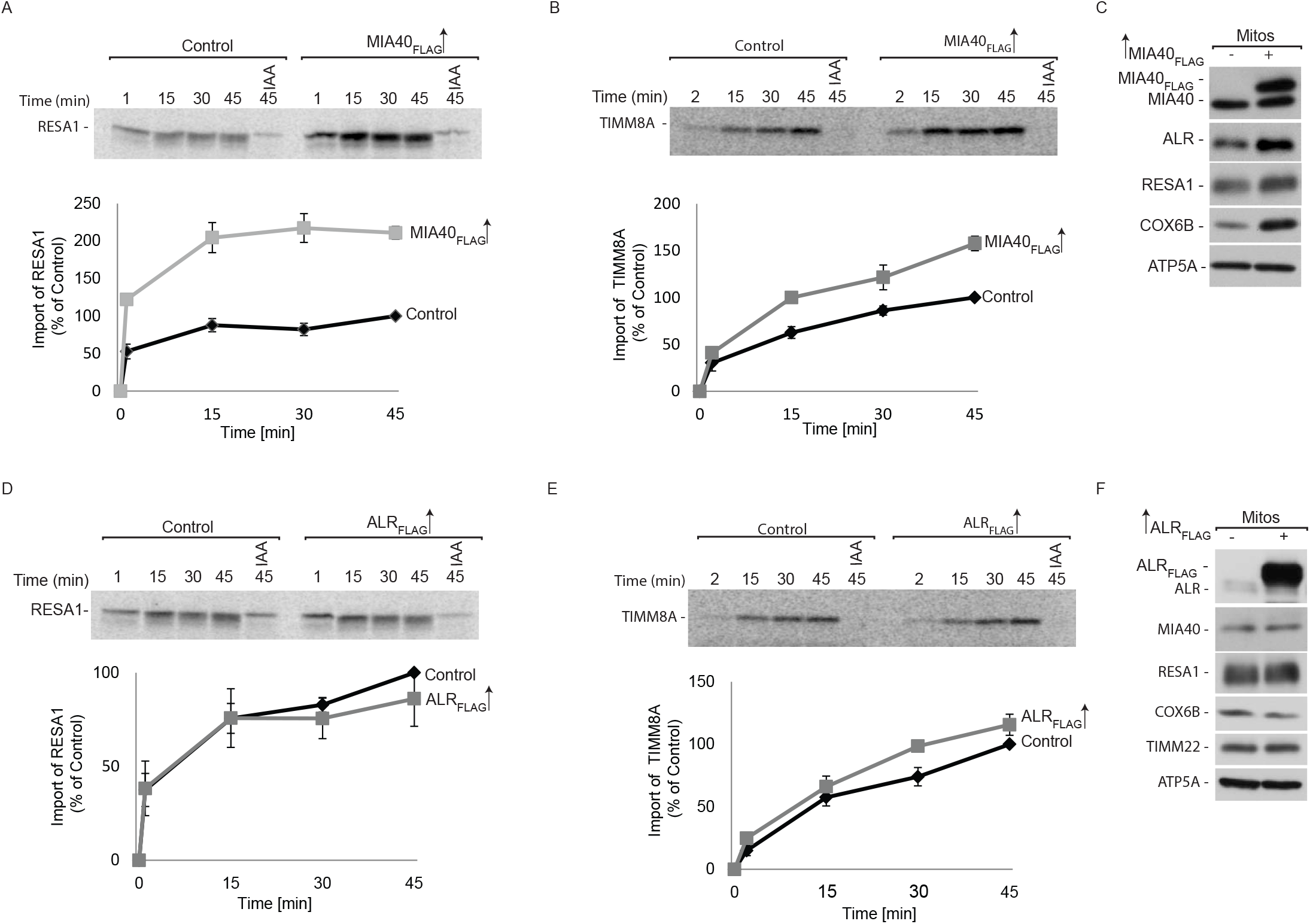
MIA40 facilitates RESA1 import into mitochondria. (A, B) Radiolabeled [^35^S]RESA1 (A) and [^35^S]TIMM8A (B) precursors were imported into mitochondria that were isolated from cells that expressed MIA40_FLAG_. The samples were analyzed by reducing SDS-PAGE and autoradiography. The results of three biological replicates were analyzed, quantified, and normalized to control mitochondria at 45 min. The data are expressed as mean ± SEM. IAA, iodoacetamide. (C) Mitochondria were isolated from Flp-In 293 T-REx cells that expressed MIA40_FLAG_ and control cells. The samples were analyzed by SDS-PAGE and Western blot. Mitos, mitochondria. (D, E) Radiolabeled [^35^S] RESA1 (D) and [^35^S] TIMM8A (E) precursors were imported into mitochondria that were isolated from Flp-In 293 T-REx cells that expressed ALR_FLAG_ and control cells. The samples were analyzed by reducing SDS-PAGE and autoradiography. The results of three biological experiments were analyzed, quantified, and normalized to control mitochondria at 45 min. The data are expressed as mean ± SEM. (F) Mitochondria were isolated from Flp-In 293 T-REx cells that expressed ALR_FLAG_ and control cells. The samples were analyzed by SDS-PAGE and Western blot.

To further substantiate the involvement of the MIA pathway in the import of RESA1, we developed a cell line with deletion of the CPC motif in only one allele of the gene (HEK293 MIA40 WT/Del^53-60^) (Fig 6A). The Western blot analysis of HEK293 MIA40 WT/Del^53-60^ cells revealed two bands that were specific to MIA40 (wildtype MIA40 and Del^53-60^ MIA40; Fig 6B). In this cell line, we observed lower levels of MIA40 substrates, including COX6B, TIMM9, and TIMM22. The levels of other mitochondrial proteins, such as MIC19 and ATP5A, remained unchanged (Fig 6B). The decrease in MIA40-dependent proteins was successfully rescued by the exogenous overexpression of MIA40 (Fig 6C, lanes 3 and 4). Thus, we established that the observed phenotype was specific to MIA40 depletion. Furthermore, we used mitochondria that were isolated from MIA40 WT/Del^53-60^ to measure the import of radiolabeled [^35^S] RESA1. The import efficiency of RESA1 was reduced to almost 50% in this mutant haploinsufficient cells when compared with wildtype cells (Fig 6D). We also observed a 40% reduction of the import of the classic MIA40 substrate TIMM8A (Fig 6E). Again, these observations indicate that RESA1 utilizes the MIA40 pathway for import into mitochondria, and it is a newly identified non-canonical substrate of MIA40.

**Figure 6.**
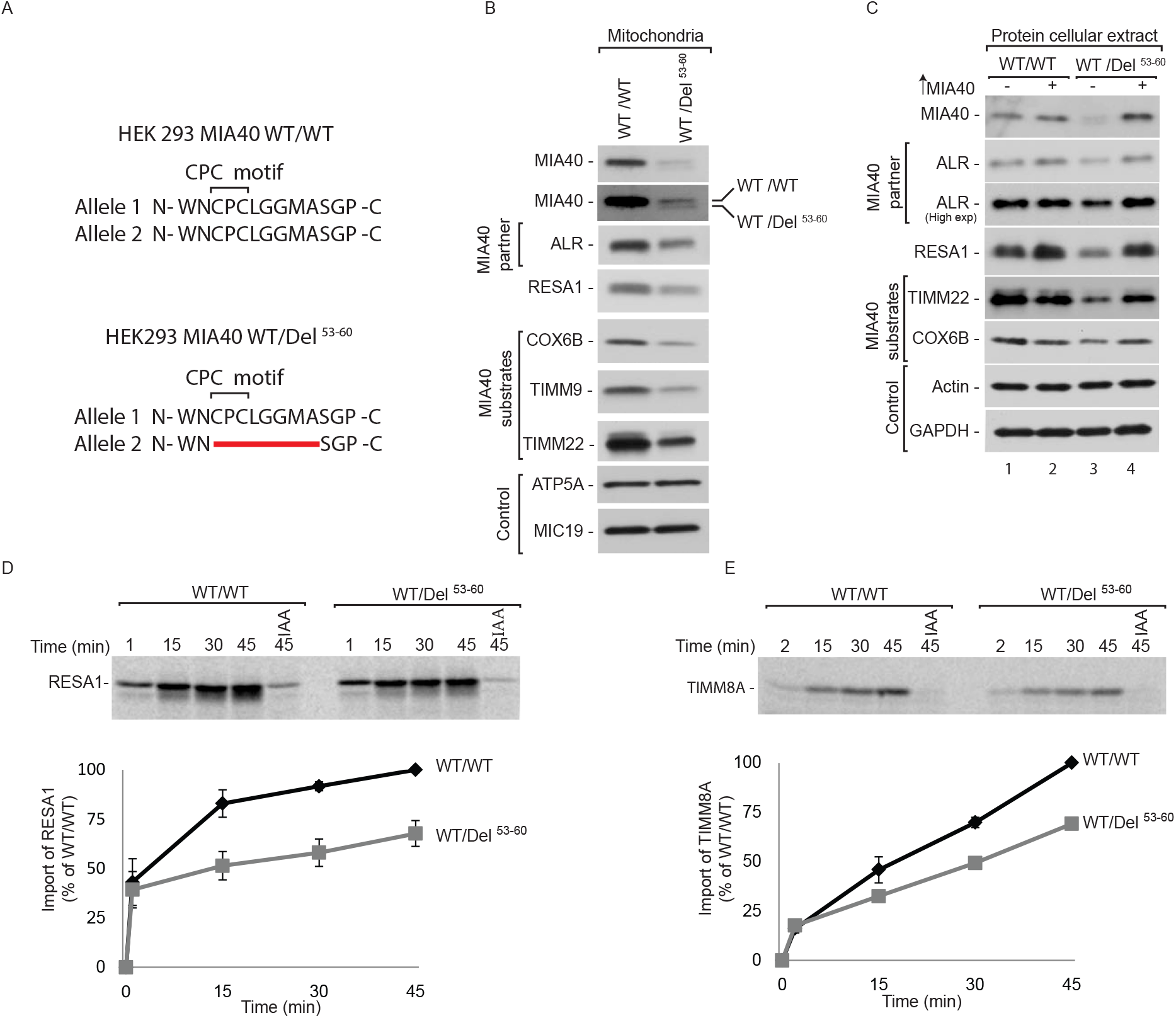
MIA40 is involved in the import and biogenesis of RESA1. (A) Schematic representation of MIA40 sequence that contains the CPC motif in HEK293 MIA40 WT/WT and HEK293 MIA40 WT/Del^53-60^ cells. (B) Cellular protein extracts were isolated from HEK293 MIA40 WT/WT and WT/Del^53-60^ cells. The samples were analyzed by SDS-PAGE and Western blot. (C) Cellular protein extracts were isolated from HEK293 MIA40 WT/WT and WT/Del^53-60^ cells that were transfected with a plasmid that encoded MIA40 or an empty vector. The samples were subjected to reducing SDS-PAGE and Western blot. (D, E) Radiolabeled [^35^S] RESA1 (D) and [^35^S] TIMM8A (E) precursors were imported into mitochondria that were isolated from HEK293 MIA40 WT/WT and WT/Del^53-60^ cells. The samples were analyzed by reducing SDS-PAGE and autoradiography. The results of three biological replicates were analyzed, quantified, and normalized to control mitochondria at 45 min. The data are expressed as mean ± SEM. IAA, iodoacetamide.

### RESA1 pathological mutants are defective in mitochondrial import

Mutations in RESA1 are associated with the defective assembly and function of respiratory chain complexes in patients diagnosed with mitochondrial encephalopathy (Higuchi et al, 2018; Martinez Lyons et al, 2016). The first reported case presented with a biallelic compound heterozygous mutation which led to two forms of RESA1: RESA1 with a single amino acid mutation (RESA1-Y137C) and RESA1 with a deletion of exon 2 (RESA1-exon2^Δ^). However these mutant proteins were undetectable in patient-derived cultured skin fibroblasts (Martinez Lyons et al, 2016). Therefore, we characterized these RESA1 mutants to elucidate the mechanism by which protein loss occurs under pathological conditions.

In order to do so, we overexpressed wildtype RESA1 and its mutant variants (RESA1-Y137C_HIS_ and RESA1-exon2^Δ^_HIS_) in HEK293 cells. We detected significantly lower levels of both variants compared with wildtype (Fig 7A), which is consistent with protein loss in fibroblasts from patients (Martinez Lyons et al, 2016). Next, we investigated the localization of mutant proteins to mitochondria and observed a proportional decrease in the mitochondrial fraction (M) together with a slight increase of the mutant proteins in the cytosolic fraction (C) (Fig 7B, lanes 3, 9 and 12). Despite only a slight defect in mitochondrial localization of mutant RESA1-Y137C_HIS_, diminished steady-state levels of protein support the possibility of degradation. The mitochondrial fraction of RESA1-Y137C_HIS_ presents IMS localization, which was confirmed by mitoplasting (Fig 7C). As a control, we observed the efficient localization of mitochondrial proteins, such as cytochrome c, ATP5A, TOMM20, and HSP60, in the mitochondrial fraction, whereas the cytosolic protein tubulin was present only in the cytosolic fraction.

**Figure 7.**
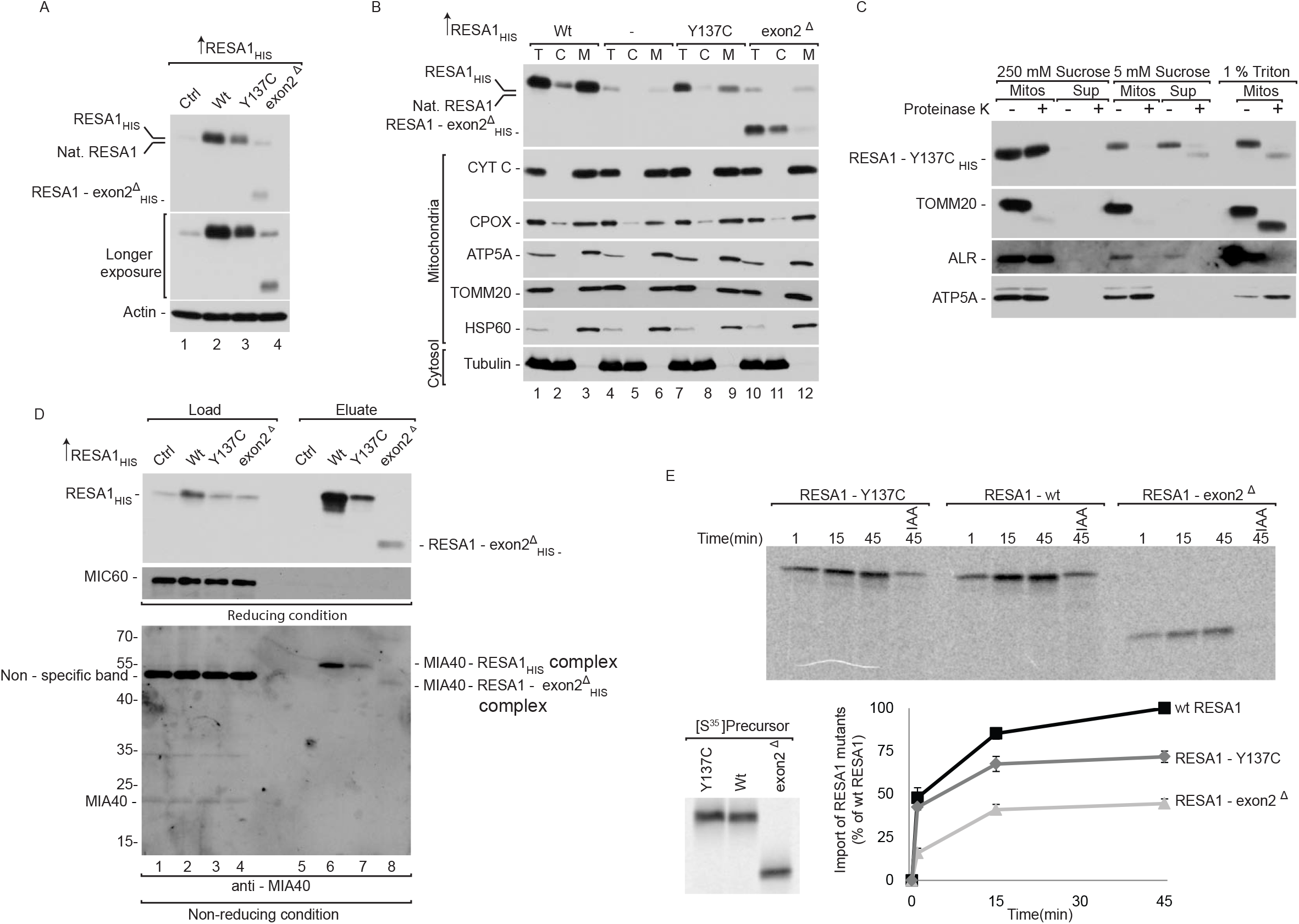
RESA1 pathological mutants are import-defective. (A) Cellular protein extracts were isolated from HEK293 cells that were transfected with a plasmid that encoded wildtype or mutant RESA1. The samples were analyzed by reducing SDS-PAGE and Western blot. (B) Cellular fractions were prepared from HEK293 cells that were transfected with a plasmid that encoded wildtype or mutant RESA1. The fractions were analyzed by reducing SDS-PAGE and Western blot. T, total; C, cytosol; M, mitochondria. (C) Localization of overexpressed RESA1-Y137C analyzed by limited degradation by proteinase K in intact mitochondria (250 mM sucrose), mitoplasts (5 mM sucrose), and mitochondrial lysates (1% Triton X-100) that were isolated from HEK293 cells. The samples were analyzed by SDS-PAGE and Western blot. Mitos, mitochondria; Sup, post-mitochondria supernatant. (D) HEK293 cells that expressed wildtype or mutant RESA1_HIS_ were solubilized, and the affinity purification of RESA1_HIS_ was performed. The samples were analyzed by reducing and non-reducing SDS-PAGE and Western blot. Load: 3%. Eluate: 100%. (E) Equal amounts of wildtype and mutant radiolabeled [^35^S]RESA1 precursors were imported into mitochondria that were isolated from HEK293 cells. The samples were analyzed by reducing SDS-PAGE and autoradiography. The results of three biological replicates were analyzed, quantified, and normalized to wildtype RESA1 at 45 min. The data are expressed as mean ± SEM. IAA, iodoacetamide.

Improperly folded mitochondrial IMS proteins can be substrates of proteases or can be retro-translocated to the cytosol (Bragoszewski et al, 2015). We performed a molecular dynamics simulation to understand the structural features that contribute to the instability of RESA1-Y137C. Homology modeling of the mutant suggested the possibility of two alternative disulfide bridges (C100-C137 and C111-C137) in addition to the native disulfide bond (C100-C111) that could potentially destabilize the structure (Fig EV5A). To further understand the dynamics of the mutant proteins, we subjected the wtRESA1 and RESA1-Y137C mutants to molecular dynamics simulations, which revealed greater flexibility in the two loop regions (Q159-K166 and D198-K202) of the RESA1-Y137C (Fig EV5B). We also identified the possibility of new salt bridge formation between lysine (K133) and aspartic acid (D136) in the RESA1-Y137C mutant (Fig EV5C). Thus, we propose that the additional cysteine at the 137th position leads to alternative disulfide bridges that compete with the native disulfide bond (C100-C111) causing misfolding and destabilization leading to structural instability, which can in turn increase the possibility of protein removal from the IMS. We previously established that RESA1 interacts with MIA40 for its efficient import into mitochondria. Therefore, we investigated the influence of these mutations on the MIA40-RESA1 interaction. Affinity purification *via* RESA1-Y137C_HIS_ and RESA1-exon2^Δ^_HIS_ revealed an uncompromised interaction with MIA40 (Fig 7D; lanes 6-8). The lower level of co-purified MIA40 could be attributed to lower expression of the mutant variants (Fig 7D, lanes 2-4). Finally, *in organello* import of the RESA1 mutants revealed a 50% decrease in import efficiency for the RESA1-Y137C mutant and a nearly 70% decrease for RESA1-exon2^Δ^ compared with wildtype (Fig 7E). Therefore, even though the mutant variants can interact with MIA40, their import into mitochondria is specifically impaired. As shown before, the steady-state levels of both mutated proteins are very low which could be the result of a combination of their intrinsic instability, low import efficiency and the inability of being productively maintained in the IMS, thus, making them a subject of degradation.

### The proteasome degrades cytosol-localized RESA1 mutants

Mitochondrial IMS proteins are efficiently degraded by the proteasome either before import or after retrotranslocation. We followed the degradation kinetics of mutant proteins by performing a cycloheximide-chase experiment (Fig 8A) and found that the mutant proteins degraded faster than wildtype (Fig 8B, lanes 3 and 5; Fig EV6A). We then evaluated the role of the proteasome in the degradation of mutant proteins by inhibiting the proteasome with MG132 (Fig 8C). Effective inhibition of the proteasome was confirmed by an increase in the ubiquitination of cellular proteins, evaluated by Western blot (Fig EV6B). We found that MG132 treatment did not rescue the degradation of already synthesized mutant proteins (Fig 8D, lanes 2 and 3). This indicates that the degradation of previously synthesized and likely mitochondrially localized protein variants occurs independently of the UPS. In contrast, we rescued the degradation of RESA1 mutants by proteasome inhibition under active translation, suggesting that the newly synthesized or cytosol-localized proteins were sensitive to proteasome-mediated degradation (Fig 8D, lanes 4 and 5). Next, we investigated whether the stabilization of mutant proteins in the cytosol increases their translocation to mitochondria (Fig 8E). Indeed, in the presence of MG132 mediated proteasome inhibition, the levels of RESA1 mutant proteins increased in the cytosol and more interestingly a larger portion of the mutant proteins was localized in mitochondria (Fig 8F, lanes 3, 7, 10, 14). Inhibition of the proteasome was confirmed by an increase in protein ubiquitination (Fig EV6C).

**Figure 8.**
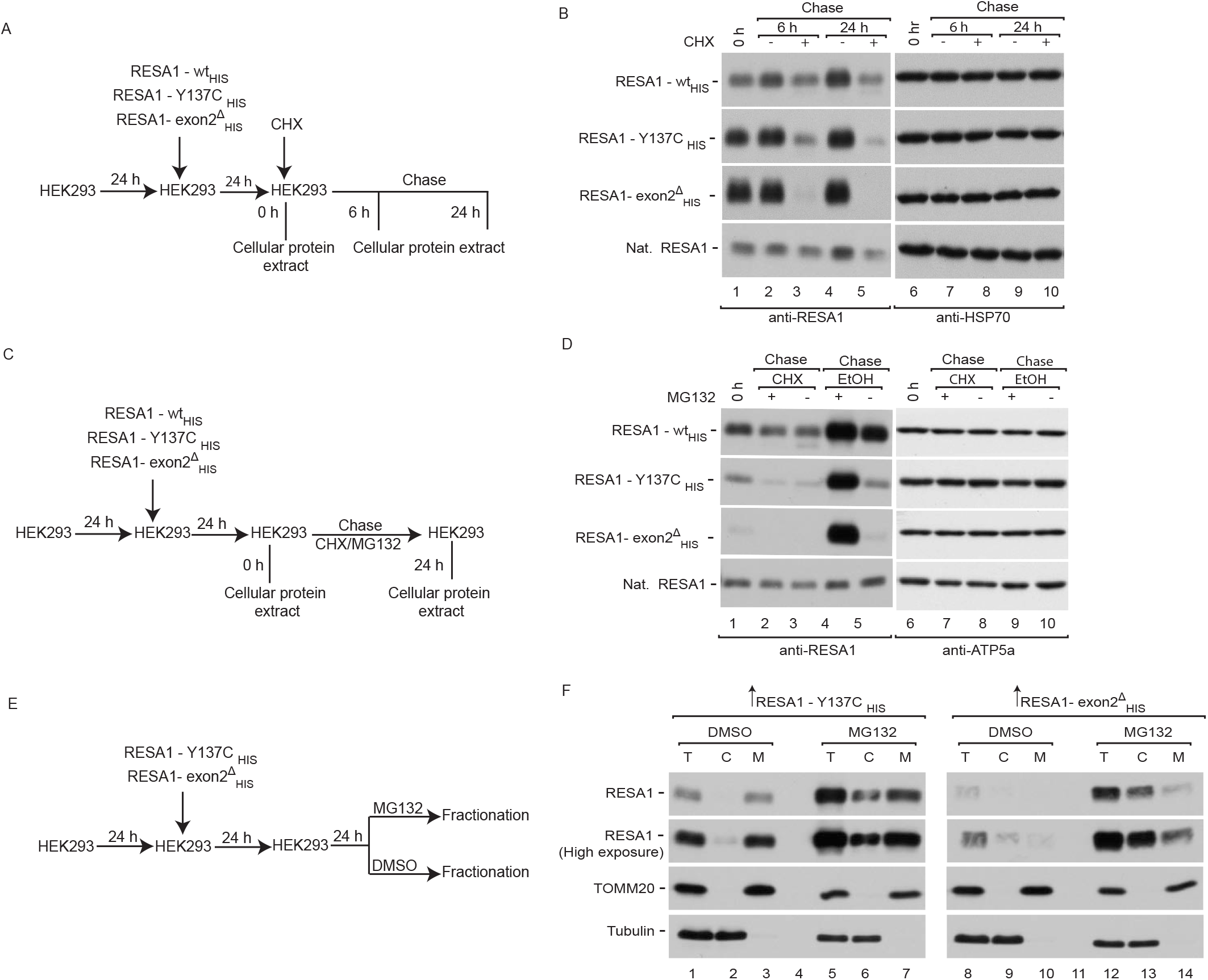
The proteasome degrades cytosol-localized RESA1 mutants. (A) Schematic representation of protein stability assay. (B) HEK293 cells that expressed wildtype or mutant RESA1_HIS_ were treated with CHX for the indicated times, and protein extracts were isolated. The samples were analyzed by reducing SDS-PAGE and Western blot. (C) Schematic representation of combined translation and proteasome inhibition assay. (D) HEK293 cells that expressed wildtype or mutant RESA1_HIS_ were treated with CHX and/or MG132 for the indicated times, and protein extracts were isolated. The samples were analyzed by reducing SDS-PAGE and Western blot. CHX, cycloheximide. (E) Schematic representation of the cellular localization assay. (F) Cellular fractions were prepared from HEK293 cells that expressed mutant RESA1_HIS_ and were treated with MG132. The samples were analyzed by reducing SDS-PAGE and Western blot.

### Rescue of RESA1 mutant proteins in a patient cell line upon proteasome inhibition

The ability to increase the levels of pathogenic variants of mitochondrial proteins prompted us to investigate whether the defects can be rescued in cell lines from patients. We inhibited proteasome machinery in immortalized fibroblasts that were derived from the subject who presented with both pathogenic variants in RESA1 (mt4229i). We used MG132 and other two clinically used proteasome inhibitors (bortezomib and carfilzomib). Proteasome inhibition increased the levels of both the mutation-carrying RESA1 proteins (RESA1-Y137C & RESA1-exon2^Δ^) in the patient cells (Fig 9A). As a control for the experiment, we also observed an increase in the levels of HSP70 upon treatment with proteasome inhibitors as reported previously (Awasthi & Wagner, 2005; Kim et al, 1999). Interestingly, upon inhibition we also observed an increase in the levels of COX6A and COX6B, the structural subunits of complex IV which are affected in patient. Further, we tested the mitochondrial localization of the rescued mutants upon proteasome inhibition and observed an increase in localization of the rescued mutants to mitochondria in the patient cell line (Fig 9B). These results together with the previous findings using HEK293 cells, substantiated the involvement of the UPS in the degradation of misfolded/mislocalized mitochondrial proteins. Thus, we confirmed the influence of the proteasome system on protein levels under pathological conditions. Since proteasome inhibition increased RESA1 levels inside mitochondria, we tested whether this effect, which involved also the mutant variants, could rescue the complex IV deficiency of patient cells (Martinez Lyons et al., 2016). Hence, we performed spectrophotometric kinetic measurements of cytochrome c oxidation. Treatment of patient fibroblasts with inhibitors of proteasome led to approximately 35% increase in complex IV activity as assessed by the oxidation of cytochrome c (Fig 9C). To further verify whether the observed significant effect was attributable to higher levels of RESA1, we combined bortezomib treatment with the overexpression of wildtype and the two different mutant versions of RESA1. Fibroblasts that expressed wtRESA1 and RESA1-Y137C exhibited a significant 50% increase in complex IV activity, and RESA1-exon2^Δ^ expression did not rescue the defective phenotype (Fig 9D). This strongly suggests that RESA1-Y137C is a functionally competent protein that can effectively modulate the activity of complex IV if allowed to reach the IMS. Thus, the defective phenotype that is observed in this patient is principally due to mislocalisation and/or premature degradation of proteins by the proteasome system, and proteasome inhibition may be beneficial for patients under similar conditions.

**Figure 9.**
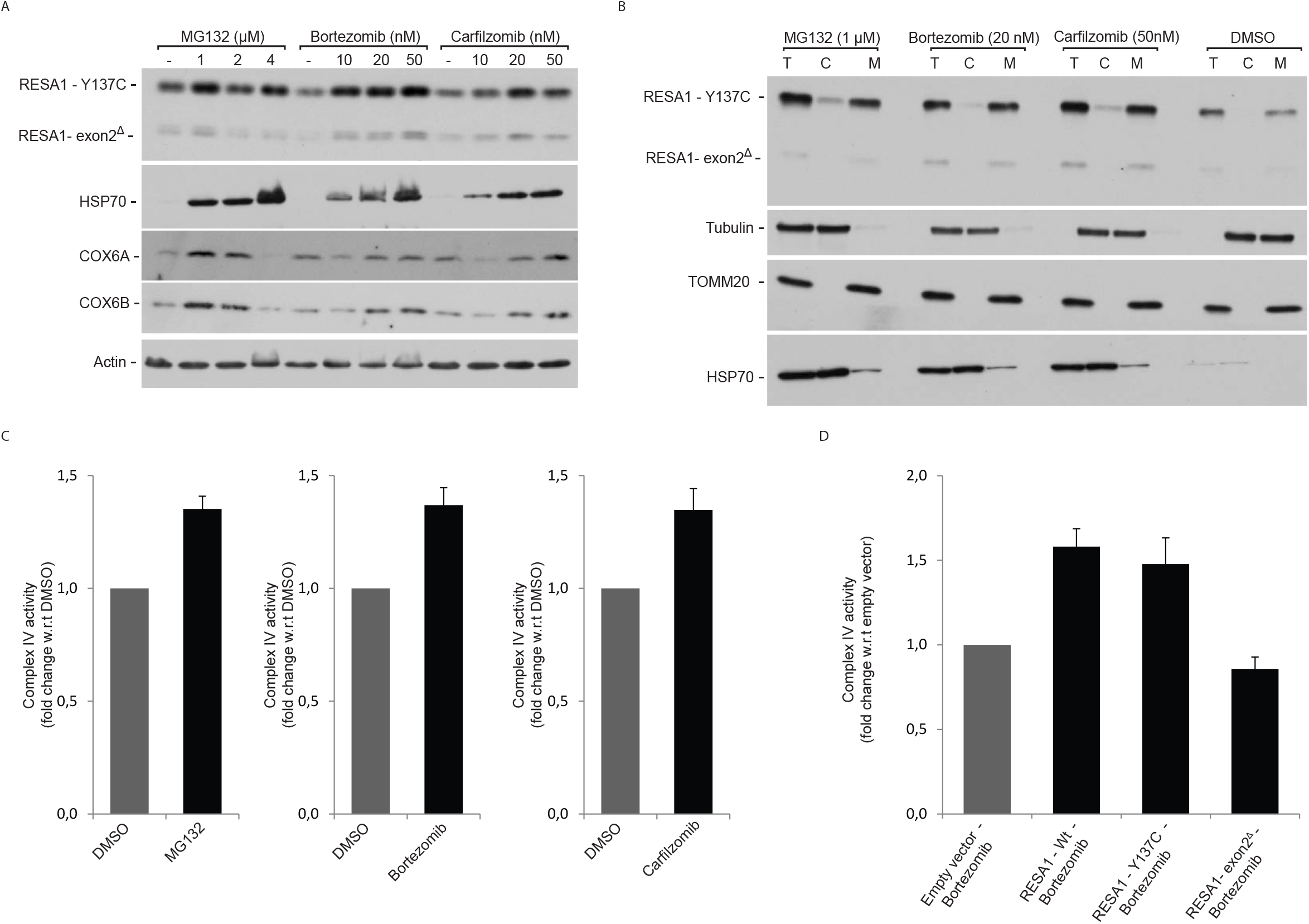
Inhibition of the proteasome rescues mitochondrial levels of RESA1 in patient-derived fibroblasts. (A) Cellular protein extracts were isolated from immortalized patient-derived skin fibroblasts that were treated with the indicated concentrations of MG132, bortezomib, or carfilzomib. The samples were analyzed by reducing SDS-PAGE and Western blot. (B) Cellular fractions were prepared from immortalized patient-derived skin fibroblasts that were treated with the indicated concentrations of MG132, bortezomib, or carfilzomib. The samples were analyzed by reducing SDS-PAGE and Western blot. T, total; C, cytosol; M, mitochondria. (C) Immortalized patient-derived skin fibroblasts were treated with MG132, bortezomib, or carfilzomib for 12 h and recovered for another 6 h. The fibroblasts were harvested, and complex IV activity was measured in digitonized cellular extracts. Activity is expressed as millimoles of oxidized cytochrome c per minute per milligram of protein. The results of three biological replicates were analyzed, quantified, and normalized to DMSO treated samples and the data are expressed as mean ± SEM. (D) Immortalized patient-derived skin fibroblasts were transfected with a plasmid that encoded wildtype or mutant RESA1 (RESA1-Y137C and RESA1-Ex2^Δ^) for 48 h, and fibroblasts were treated with bortezomib during the last 12 h. The fibroblasts were harvested, and complex IV activity was measured in digitonized cellular extracts. Activity is expressed as millimoles of oxidized cytochrome c per minute per milligram of protein. The results of three biological replicates were analyzed, quantified, and normalized to empty vector-bortezomib treated samples and the data are expressed as mean ± SEM.

## Discussion

The present study established a recently identified IMS protein, RESA1, as a non-canonical substrate of MIA40. RESA1 exists as an oxidized protein in the IMS of mitochondria, and MIA40 facilitates its import into IMS. Whole-exon sequencing of a patient who was diagnosed with leukoencephalopathy recently revealed biallelic heterozygous mutations of RESA1 that led to absence of the protein (Martinez Lyons et al, 2016). We discovered that the proteasome-mediated degradation of pathogenic RESA1 mutant proteins accounted for nearly the complete absence of protein. When cells expressing both point mutation and deletion instable RESA1 proteins were treated with proteasome inhibitors, the degradation of the mutated proteins was prevented and their localization to mitochondria was increased. In the patient-derived cells this increase in RESA1 levels was associated with an increase in complex IV activity.

Most mitochondrial proteins are translated in the cytosol where they are inevitably in proximity to the proteasome, which is the main cytosolic protein quality control machinery. Although mitochondria contain autonomous protein homeostasis mechanisms, there are many examples of mitochondrial proteins that are degraded by the proteasome. Integral proteins of the OMM, such as MCL1, MFN1, BCL2, FIS1, and DRP1, can undergo degradation in the cytosol by the proteasome (Azad et al, 2006; Karbowski et al, 2007; Neutzner et al, 2008; Yonashiro et al, 2006; Zhong et al, 2005; Ziviani et al, 2010). Similarly, the proteasome regulates the turnover of certain IMS proteins, such as apo-cytochrome c and endonuclease G (Pearce & Sherman, 1997; Radke et al, 2008). We previously demonstrated that proteasome machinery regulates the mitochondrial IMS proteome through the degradation of misfolded or mislocalized proteins in both yeast and mammalian systems (Bragoszewski et al, 2013; Bragoszewski et al, 2015). Interestingly, Bragoszewski et al., showed that IMS proteins, such as COX6B, can follow a retrograde route of transport to the cytosol for proteasome-mediated degradation. The efficient release of this class of proteins from mitochondria is facilitated by the reduction of intramolecular disulfide bridges and unfolding, which allows for translocation through the TOM channel (Bragoszewski et al, 2015).

The interplay between mitochondria and the proteasome extends beyond the routine turnover of mitochondrial proteins. Upon the failure of mitochondrial import, excess mitochondrial precursor proteins in the cytosol is eliminated by the proteasome mediated degradation prior to their import (Bragoszewski et al, 2013; Wrobel et al, 2015). Moreover, substrates of the MIA pathway in yeast were found to be ubiquitinated and accumulated in response to proteasome inhibition, even in the presence of active import machinery (Bragoszewski et al, 2013; Kowalski et al, 2018). Thus, the proteasome system acts as a vital checkpoint for the improper localization of proteins and promotes efficient mitochondrial IMS protein biogenesis (Bragoszewski et al, 2013; Bragoszewski et al, 2017).

The present data demonstrate that the degradation of mitochondrial proteins by the proteasome is linked to their synthesis, in which overexpressed RESA1 mutant variants are stabilized by the proteasome inhibitor MG132 only in the absence of the translation inhibitor cycloheximide. This conclusion is supported by previous findings that a large fraction of newly synthesized proteins is subjected to proteasome-mediated degradation (Schubert et al, 2000), and mitochondrial proteins are co-ubiquitinated during translation (Duttler et al, 2013; Wang et al, 2013). This is consistent with our findings in which mislocalized immature RESA1 mutant variants were degraded by the proteasome. Interestingly, mutant variants of RESA1 interact with MIA40, raising the possibility that they follow retro-translocate-mediated degradation through inefficient folding and accumulation in mitochondria.

Therefore, we cannot completely exclude the involvement of mitochondrion-localized proteases in the turnover of these mutant proteins. These findings indicate that the proteasome and mitochondrial import machinery compete for the same pool of mitochondrial proteins to promote efficient mitochondrial IMS protein biogenesis and quality control. Thus, the proteasome plays an important and previously unexpected role in the control and maintenance of mitochondrial proteome homeostasis.

In the present study, we characterized two pathogenic mutants of RESA1 that are associated with mitochondrial leukoencephalopathy. These proteins were less efficiently imported into mitochondria and degraded faster than wild type protein. The faster rate of degradation of mutant proteins is corroborated by the apparent lack of RESA1 in patient cells (Martinez Lyons et al, 2016). Inhibition of the proteasome rescued mutant protein levels and increased their mitochondrial levels, accompanied by higher levels of complex IV subunits (e.g., COX6B and COX6A) and an increase in the activity of complex IV. The recovery of complex IV activity was also achieved by the overexpression of RESA1-Y137C protein, suggesting that the mutant protein, if allowed to be imported into mitochondria, was sufficiently active to ameliorate the pathological phenotype. These data strongly suggest that the pathological conditions that are observed in patients with mutant RESA1 can result from mutant mislocalization, thus leading to excessive degradation by the proteasome rather than excessive degradation from the loss of function of mutant proteins. Analogously, other studies reported that the overexpression of mutant versions of mitochondrial proteins, such as NUBPL and FOXRED1, was able to reverse the pathological phenotype in patient cells (Formosa et al, 2015; Tucker et al, 2012). The stimulation of mitochondrial import of the pathogenic mutant CHCHD10 was recently proposed as a possible therapeutic strategy for amyotrophic lateral sclerosis (Lehmer et al, 2018). Our data suggest that the inhibition of excessive degradation by the proteasome can rescue mitochondrial function in diseases that are associated with the mislocalization and premature degradation of mitochondrial proteins.

Additionally, pathogenic mutations of various mitochondrial proteins (e.g., NDUFAF3, NDUFAF4, FOXRED1, COA5, COA6, COX6B, CHCHD10, and tafazzin) were associated with the secondary loss of other mitochondrial proteins, which can aggravate the disease (Dudek et al, 2013; Modjtahedi & Kroemer, 2016; Nouws et al, 2012). In patient-derived fibroblasts, structural subunits of cytochrome c oxidase COX6A and COX6B decreased in parallel with RESA1 and were rescued by proteasome inhibition. Thus, proteasome inhibition prevented also a secondary loss of mitochondrial proteins. Importantly, mitochondrial biogenesis has been previously shown to be a rescue mechanism for defective mitochondria (Hansson et al, 2004; Kuhl et al, 2017).

Mitochondrial diseases affect approximately one in 2000 people and may arise at any age with a wide range of clinical symptoms (Gorman et al, 2016; Suomalainen & Battersby, 2018). Among these diseases are several genetic mitochondrial pathologies that are associated with the apparent loss of mutated proteins that are often associated with the more generalized deficiency of mitochondria protein biogenesis (Modjtahedi & Kroemer, 2016). Despite dramatic improvements in the genetic and metabolic diagnoses of these severe progressive diseases, no curative treatments have been discovered. The current treatment regimen for respiratory chain deficiency is mostly metabolite supplementation with CoQ10, creatinine, or riboflavin, which is only a symptomatic treatment and does not ameliorate the underlying pathological condition. In the present study, we used bortezomib and carfilzomib (i.e., two proteasome inhibitors that have been approved for the treatment of patients with multiple myeloma and mantle cell lymphoma) to reverse the pathological phenotype in fibroblasts that were derived from a patient who suffered from mitochondrial leukoencephalopathy. Clinically approved proteasome inhibitors, such as bortezomib, may be applied as a therapeutic intervention to treat mitochondrial diseases that are associated with the excessive degradation of mitochondrial proteins by the proteasome that is primarily attributable to their mutation but also attributable to defective import and biogenesis in mitochondria.

## Materials and Methods

### Cell lines and growth conditions

HeLa, HEK293, and Flp-In T-REx 293 cells were cultured at 37°C with 5% CO_2_ in standard Dulbecco’s Modified Eagle Medium (DMEM) supplemented with 10% (v/v) heat-inactivated fetal bovine serum, 2 mM L-glutamine, 100 U/ml penicillin, and 10 μg/ml streptomycin sulfate. The cells were grown in DMEM that contained low glucose (1.1 g/L) or galactose (1.8 g/L) as described below. Immortalized skin fibroblast cell lines (Ai and mt4229i) were grown in standard DMEM supplemented with 10% (v/v) heat-inactivated fetal bovine serum, 2 mM L-glutamine, 100 U/ml penicillin, 10 μg/ml streptomycin sulfate, 1 mM sodium pyruvate, and 50 μg/ml uridine. Fibroblast cells were grown in DMEM that contained low glucose (1.1 g/L) or galactose (1.8 g/L) as described below. The MIA40 deletion cell line (HEK293 MIA40 WT/Del Del^53-60^) was generated using guide RNA that targeted exon 3 of MIA40 ORF cloned into the pX330 vector (Addgene, catalog no. 42230). The recombinant plasmid was co-transfected into HEK293 cells with green fluorescent protein (GFP)-expressing plasmid. Single cells that expressed GFP were sorted into each well of the 96-well plate and grown at 37°C under 5% CO_2_. Deletion that corresponded to amino acids 53-60 of MIA40 was confirmed by sequencing.

### siRNA and transfection

For gene knockdown, the following mRNA sequences were targeted: MIA40_1 (5’-ATAGCACGGAGGAGATCAA-3’), MIA40_2 (5’-GGAATGCATGCAGAAATAC-3’), ALR_1 (5’-GGAGTGTGCTGAAGACCTA-3’), ALR_2 (5’-GCATGCTTCACACAGTGGCTGT-3’), RESA1_1 (5’-GATGGTGTTGATAAGGATGA-3’), and RESA1_2 (5’-GGCACATGATGGACAGGTTAA-3’). The transfection of HEK293 cells was performed in low-glucose DMEM using oligofectamine as per manufacturer’s instructions. After 24 h in low glucose, the medium was changed to galactose-containing DMEM for the next 48 h and cells were harvested. As a control, the cells were transfected with Mission siRNA universal negative control (SIC001, Sigma).

### Cellular protein lysate

Cells were harvested and lysed in radioimmunoprecipitation assay (RIPA) buffer (65 mM Tris-HCl [pH 7.4], 150 mM NaCl, 1% v/v NP 40, 0.25% sodium deoxycholate, 1 mM ethylenediaminetetraacetic acid [EDTA], and 2 mM phenylmethylsulfonyl fluoride [PMSF]) for 30 min at 4°C. The lysate was clarified by centrifugation at 14,000 × *g* for 30 min at 4°C. The supernatant was collected, and the protein concentration was measured by the bicinchoninic acid protein assay (ThermoFischer Scientific). The supernatant was diluted in Laemmli buffer that contained either 50 mM DTT or 50 mM IAA for reducing and nonreducing conditions, respectively.

### FLAG-tag affinity purification

A total of 4 × 10^6^ cells were seeded and grown in low-glucose medium for 24 h and then induced with tetracycline (100 ng/ml) in galactose medium for 24 h. The cells were harvested and solubilized in lysis buffer (50 mM Tris-HCl [pH 7.4], 150 mM NaCl, 10% glycerol, 1 mM EDTA, and 1% digitonin) supplemented with 2 mM PMSF and 50 mM IAA for 20 min at 4°C. The lysate was clarified by centrifugation at 20,000 × *g* for 15 min, and the supernatant was incubated with anti-FLAG M2 affinity gel (Sigma) for 2 h at 4°C with mild rotation. After binding, the resin was washed three times with lysis buffer without digitonin. The column-bound proteins were eluted with either 3X FLAG peptide for mass spectrometry or Laemmli buffer under reducing or non-reducing conditions for SDS-PAGE and Western blot.

### HIS-tag affinity purification

HEK293 cells were grown in low-glucose DMEM for 24 h. The cells were then transferred to DMEM and transfected with pcDNA3.1(+)-RESA1_HIS_ using Gene Juice Transfection Reagent (Merck Millipore) according to the manufacturer’s protocol. The cells were harvested 24 h after transfection and solubilized in ice-cold digitonin buffer (1% digitonin, 10% glycerol, 20 mM Tris-HCl [pH 7.4], 100 mM NaCl, and 20 mM imidazole [pH 7.4]) supplemented with 2 mM PMSF and 50 mM IAA for 20 min at 4°C. The lysate was clarified by centrifugation at 20,000 × *g* for 15 min at 4°C. The supernatant was then incubated with Ni-NTA agarose beads (Invitrogen) with mild rotation for 1.5 h at 4°C. After binding, the beads were washed three times with wash buffer (20 mM Tris HCl [pH 7.4], 100 mM NaCl, and 35 mM imidazole [pH 7.4]). Bead-bound proteins were eluted with Laemmli buffer that contained either 50 mM DTT or 50 mM IAA for reducing and non-reducing conditions, respectively. The samples were denatured and subjected to SDS-PAGE and Western blot.

### Complex IV activity

Approximately 10 × 10^6^ immortalized fibroblast cells were cultured for 24 h and subjected to proteasome inhibition. The cells were then harvested by trypsinization and resuspended in buffer A (20 mM MOPS/KOH [pH 7.4] and 250 mM sucrose) followed by digitonin treatment (0.2 mg/ml) for 5 min at 4°C. The cell lysate was centrifuged at 5,000 × *g* for 3 min. The supernatant that contained the cytosolic fraction was discarded, and the pellet was resuspended in buffer B (20 mM MOPS/KOH [pH 7.4], 250 mM sucrose, and 1 mM EDTA). The solution was centrifuged at 10,000 × *g* for 3 min, and the pellet that contained mitochondria was resuspended in 10 mM potassium phosphate buffer (pH 7.4) and frozen-thawed three times in liquid nitrogen immediately before starting the spectrophotometric assays (Tiranti et al, 1995). Complex IV activity was measured as a decrease in absorbance at 550 nm that was caused by the oxidation of cytochrome c. Activity is expressed as nanomoles of cytochrome c per minute per milligram of protein.

### Mitochondrial isolation and fractionation

HEK293 cells were harvested and resuspended in ice-cold trehalose buffer (10 mM HEPES-KOH [pH 7.7], 300 mM trehalose, 10 mM KCl, 1 mM EDTA, and 1 mM ethylene glycol-bis[β-aminoethyl ether]-N,N,N’,N’-tetraacetic acid [EGTA]) supplemented with 2 mg/ml bovine serum albumin (BSA) and 2 mM PMSF. After homogenization in a Dounce glass homogenizer (Sartorius), the homogenate was clarified by centrifugation at 1000 × *g* for 10 min at 4°C. The supernatant was centrifuged at 10,000 × *g* for 10 min at 4°C. The pellet was resuspended in trehalose buffer without BSA, and protein levels were quantified using the Bradford assay.

Human fibroblasts were harvested and resuspended in ice-cold isotonic buffer (10 mM MOPS [pH 7.2], 75 mM mannitol, 225 mM sucrose, and 1 mM EGTA) supplemented with 2 mg/ml BSA and 2 mM PMSF and subjected to centrifugation at 1000 × *g* for 5 min at 4°C). The cell pellet was then resuspended in cold hypotonic buffer (10 mM MOPS [pH 7.2], 100 mM sucrose, and 1 mM EGTA) and incubated on ice for 5-7 min. The cell suspension was homogenized in a Dounce glass homogenizer (Sartorius). Cold hypertonic buffer (1.25 M sucrose and 10 mM MOPS [pH 7.2]) was added to the cell homogenate (1. 1 ml/g of cells).

The homogenate was subjected to centrifugation at 1000 × g for 10 min at 4°C to pellet the cellular debris. The supernatant that contained mitochondria was then carefully aspirated and centrifuged again. The supernatant was then subjected to high-speed centrifugation at 10000 × *g* for 10 min at 4°C to pellet mitochondria. The pellet was resuspended in isotonic buffer without BSA and quantified using the Bradford assay. Mitochondria were denatured in 2× Laemmli buffer with 50 mM DTT or 50 mM IAA. The samples were separated by SDS-PAGE and analyzed by Western blot.

For cell fractionation, after homogenization, the supernatant was then divided into three equal aliquots. One aliquot was saved as the total fraction. Two aliquots were centrifuged at 10,000 × *g* for 10 min at 4°C to collect the pellet and supernatant for the mitochondrial and cytosolic fractions, respectively. The proteins from the total and cytosolic fractions were precipitated by pyrogallol red precipitation. Finally, all three fractions were denatured in urea sample buffer with 50 mM DTT for SDS-PAGE and Western blot.

### Radioactive precursor synthesis and in organello import

The cDNA of precursors was cloned into a pTNT vector under the SP6 promoter, and radiolabeled precursors were synthesized using [^35^S]methionine and the TNT SP6 Quick Coupled Transcription/Translation system (Promega). The precursors were precipitated by ammonium sulfate and reduced in urea buffer (6 M urea and 60 mM MOPS-KOH [pH 7.2]) with 50 mM DTT. The import of radiolabeled precursors into isolated mitochondria was performed at 30°C in import buffer (250 mM sucrose, 80 mM potassium acetate, 5 mM magnesium acetate, 5 mM methionine, 10 mM sodium succinate, 5 nM adenosine triphosphate, and 20 mM HEPES/KOH [pH 7.4]). Import was stopped by the addition of 50 mM IAA and incubation on ice. Non-imported precursors were removed by proteinase K treatment for 15 min. Proteinase K was then inactivated by the addition of 2 mM PMSF, and mitochondria were washed with high-sucrose buffer (500 mM sucrose and 20 mM HEPES/KOH [pH 7.4]). The samples were solubilized in Laemmli sample buffer with 50 mM DTT and 0.2 mM PMSF and separated by SDS-PAGE and autoradiography. The efficiency of protein import was quantified by densitometry of the autoradiography images using the ImageQuantTL program. The level of import into control mitochondria at the indicated time point was set to 100%.

### Redox state analysis of proteins

The oxidation state of RESA1 was determined by direct and indirect thiol trapping assays with isolated mitochondria. For the direct thiol trapping assay, mitochondria were solubilized in Laemmli sample buffer either with 50 mM DTT or, 10 mM IAA, or 15 mM AMS (Life Technologies). For the indirect thiol trapping assay, mitochondria were treated with IAA to block free cysteine residues for 10 min at 30°C. The solution was centrifuged at 20000 × *g* for 10 min at 4°C, and mitochondria were resuspended in trehalose buffer that contained 50 mM DTT at 65°C for 15 min to reduce the disulfide bonds. Finally, mitochondria were solubilized in Laemmli buffer with AMS for 30 min at 30°C. The samples were denatured and analyzed by SDS-PAGE and Western blot.

### Mitoplasting

Isolated mitochondria were incubated on ice for 30 min in either sucrose buffer (250 mM sucrose and 20 mM HEPES/KOH [pH 7.4]) or swelling buffer (25 mM sucrose and 20 mM HEPES/KOH [pH 7.4]). The solution was divided into two equal parts. One part was treated with proteinase K (25 μg/ml) for 5 min on ice and another part was left untreated. Proteinase K was inactivated by the addition of 2 mM PMSF. Mitochondria were pelleted by centrifugation, and the supernatant fraction was subjected to pyrogallol red precipitation to recover the proteins. The pellet was then denatured in urea sample buffer with 50 mM DTT and analyzed by SDS-PAGE and Western blot.

### Immunofluorescence

HeLa cells or HeLa cells that stably expressed RESA1-HA were transfected with different subcompartment markers, namely matrix targeted photoactivatable GFP (mtPAGFP) [matrix], COX8A-DsRed [IM], and TOMM20-DsRed [OM]) and fixed with 3.7% formaldehyde after 24 h. The cells were permeabilized (1% Triton and 0.1% sodium deoxycholate in phosphate-buffered saline [PBS]) and immunostained in blocking buffer (5% goat serum and 0.1% IgG-free BSA in PBS) with anti-TOMM20 (OM; Santa Cruz Biotechnology), anti-SMAC/DIABLO (IMS; Abcam), anti-SDHA (IM; Abcam), anti-ACONITASE 2 (matrix; Abcam), anti-MDH2 (matrix; Atlas Antibodies), anti-RESA1 (Atlas Antibodies), or anti-HA (Cell Signaling Technology) antibody and the respective secondary antibody (488 or 568 Alexa Fluor, H+L- or IgG-specific, Invitrogen) and mounted with Prolong Diamond (Invitrogen).

### Super-resolution imaging and analysis

HeLa cells were transfected with different subcompartment markers (mtPAGFP [matrix], COX8A-DsRed [IM], and TOMM20-DsRed [OM]) and fixed with 3.7% formaldehyde after 24 h. The cells were permeabilized with 1% Triton and 0.1% sodium deoxycholate in PBS and immunostained in blocking buffer (5% goat serum and 0.1% IgG-free BSA in PBS) with anti-TOMM20 (Santa Cruz Biotechnology), anti-Smac/Diablo (Abcam), anti-SDHA (Abcam), anti-Aconitase-2 (Abcam), anti-MDH2 (Atlas Antibodies), and the respective secondary antibody (488 or 568 Alexa Fluor, H+L-or IgG-specific; Invitrogen) and mounted with Prolong Diamond. Acquisition was performed using an N-SIM microscope system (Nikon) equipped with a super-resolution Apo TIRF 100× 1.49 NA objective and a DU897 Ixon camera (Andor Technologies). Three-dimensional SIM image stacks were acquired with a Z-distance of 0.15 μm. All of the raw images were computationally reconstructed using the reconstruction slice system from NIS-Elements software (Nikon) while keeping the same parameters. The co-localization analysis was performed using Imaris 9.0 XT software (Bitplane Scientific Software, St. Paul, MN, USA. For each image, the threshold was applied in the same way as the controls, and Pearson’s coefficient in the co-localized volume was calculated for each image. The statistical analysis was performed using GraphPad Prism software and one-way analysis of variance (ANOVA) followed by Tukey’s multiple-comparison *post hoc* test.

### Mass spectrometry

Liquid chromatography-mass spectrometry (LC-MS) was performed at the Mass Spectrometry Laboratory (Institute of Biochemistry and Biophysics, Polish Academy of Sciences, Warsaw, Poland). The affinity-purified eluate fraction was resuspended in ammonium bicarbonate buffer (100 mM). The proteins were initially reduced by 100 mM DTT for 30 min at 57°C and alkylated with 55 mM IAA for 40 min at 57°C in the dark. The proteins were then digested overnight with 10 ng/ml trypsin at 37°C. After digestion, 0.1% trifluoroacetic acid (TFA) was added to inactivate trypsin, and the peptide suspension was clarified by centrifugation at 14000 × *g* for 20 min at 4°C. The resulting peptide mixtures were applied to an RP-18 pre-column (Waters, Milford, MA, USA) using water that contained 0.1% TFA as the mobile phase and then transferred to a nano-high-performance liquid chromatography RP-18 column (75 μM internal diameter, Waters, Milford, MA, USA) using an acetonitrile (ACN) gradient (0-35% ACN in 160 min) in the presence of 0.1% TFA at a flow rate of 250 nl/min. The column outlet was coupled directly to the ion source of an Orbitrap Elite or Q Exactive mass spectrometer (Thermo Electron, San Jose, CA, USA) working in the regime of the data-dependent MS-to-MS/MS switch. A blank run to ensure the absence of cross-contamination from previous samples preceded each analysis. The mass spectrometer was operated in the data-dependent MS2 mode. Data were acquired in a m/z range of 300-2000 Da. The Max-Quant 1.5.7.4 platform was used as described previously (Cox and Mann, 2008) using the UniProt database as a reference (accessed November 20, 2015). The Mascot search parameters were the following: mass tolerance for peptides (5 ppm), mass tolerance for fragments (0.01 Da), enzyme (trypsin), missed cleavages (1), variable modifications (oxidation [M], carbamidomethyl [C], methylthio [C], carbamidomethyl [N-term]). Label-Free-Quantification (LFQ) intensity values were calculated using the MaxLFQ algorithm as described previously (Cox et al., 2014).

### Molecular modeling

The structure of RESA1 (UniProt id: Q96BR5) was modeled using the Yasara Structure 17.1.28 package based on three Protein Data Bank (PDB) structures (1OUV, 4BWR, and 1KLX) as templates (Lüthy et al., 2002, 2004; Urosev et al., 2013). These PDB structures were selected automatically based on the combination of the blast E-value, sequence coverage, and structure quality. For each template, up to five alternate alignments with the target sequence were used, and up to 50 different conformations were tested for each modeled loop. The resulting models were evaluated according to structural quality (dihedral distribution, backbone, and side-chain packing). The model with the highest score of these that covered the largest part of the target sequence was used as the template for a hybrid model that was further iteratively improved with the best fragments (e.g., loops) that were identified among the highly scored single-template models.

### Molecular dynamics

Molecular dynamics analysis was performed using the Yasara structure with a standard Yasara2 field (Krieger et al., 2009). The initial structures were solvated in cubic boxes of water molecules, the dimensions of which allowed at least 20 Å separation/distance between the protein and box border. The general fold was initially preserved by weak distance constraints that were applied for all H-bonds that were identified in the helical regions. The molecular dynamics simulations in the isothermal-isobaric ensemble (NPT) were performed for an initial 5 ns with fixed backbone atoms (N, Cα, and C) to enable optimization of the side chains. During the next 25 ns, the system was released to evolve, and molecular dynamics snapshots were taken every 10 ps.

The conformation and flexibility of all non-helical regions were analyzed by the time evolution of phi and psi backbone angles, and are presented as Ramachandran plots. All of these analyses were performed with the R 3.3.0 package (www.r-project.org) using custom-made scripts.

## Acknowledgements

This work was financed by the National Science Centre, Poland (NCN; grant no. NCN 2012/05/B/NZ3/00781 and NCN 2015/19/B/NZ3/03272) and Ministerial funds for science within the Ideas Plus program (000263 in 2014-2017). The biobank “Cell Line and DNA Bank of Genetic Movement Disorders and Mitochondrial Diseases,” a member of the Telethon Network of Genetic Biobanks (project no. GTB12001), funded by Telethon Italy, and the EuroBioBank Network provided the control and patient-derived fibroblast specimens. We would like to thank Anabel Martinez Lyons, MRC Mitochondrial Biology Unit, University of Cambridge, who provided us with the immortalized patient fibroblasts.

## Author Contributions

KM, MW, and AC designed the study. KM performed most of the experiments and evaluated the data together with MW and AC. PS created the MIA40 deletion mutant cell line. MW created cell lines with inducible MIA40 and its mutants. MD and SD created the cell line with the inducible expression of ALR. JP performed the molecular modeling and simulation studies. DC and MD performed the mass spectrometry analysis. CB performed the microscopy experiments. EFV provided reagents and methods. PR and MZ commented on the manuscript. KM prepared the figures. KM, MW, and AC interpreted the results and wrote the manuscript. All of the authors commented on the manuscript.

## The paper explained

### Problem

Mitochondrial diseases are a clinically heterogeneous group of genetic disorders that are characterized by dysfunctional mitochondria. To date, the treatment for such diseases consists mostly of metabolite supplementation, which is only a symptomatic treatment and does not treat the underlying pathological condition.

### Results

The present study characterized a protein, RESA1, previously shown to be involved in assembly and function of respiratory chain complexes, as an intermembrane space protein and a new substrate of MIA40. We also characterized pathogenic disease-causing RESA1 mutants and found that they were imported to mitochondria more slowly than wildtype RESA1. Consequently, both these mutant proteins were mislocalized to the cytosol and degraded by the proteasome. Proteasome inhibition resulted in an increased mitochondrial localization of mutant proteins, rescuing mitochondrial complex IV deficiency in RESA1 mutated patient fibroblast. We further analyzed the fate of RESA1 mutants in immortalized patient fibroblasts. We confirmed that inhibition of the proteasome led to an increase in the mitochondrial localization of mutant proteins and rescued mitochondrial levels of other affected mitochondrial proteins. We also found that inhibition of the proteasome rescued a defect of respiratory complex IV in patient fibroblasts.

### Impact

The present study identified an important role for the proteasome in the degradation of mutant mitochondrial proteins under pathological conditions. Inhibition of the proteasome may be beneficial for patients who suffer from mitochondrial diseases that are characterized by a lower amount of mutant mitochondrial proteins. We raise the possibility that proteasome inhibitors (e.g., bortezomib and carfilzomib) that are already used clinically for cancer therapy can restore levels of mitochondrial proteins that despite being mutated, conserve some functionality when allowed to be targeted to the right compartment. With currently limited effective therapeutic options for treating mitochondrial diseases, this strategy may be promising for mitochondrial diseases, in which mitochondrial protein depletion is observed that contributes to lowering respiratory complex activity.

---

**Figure EV1.**
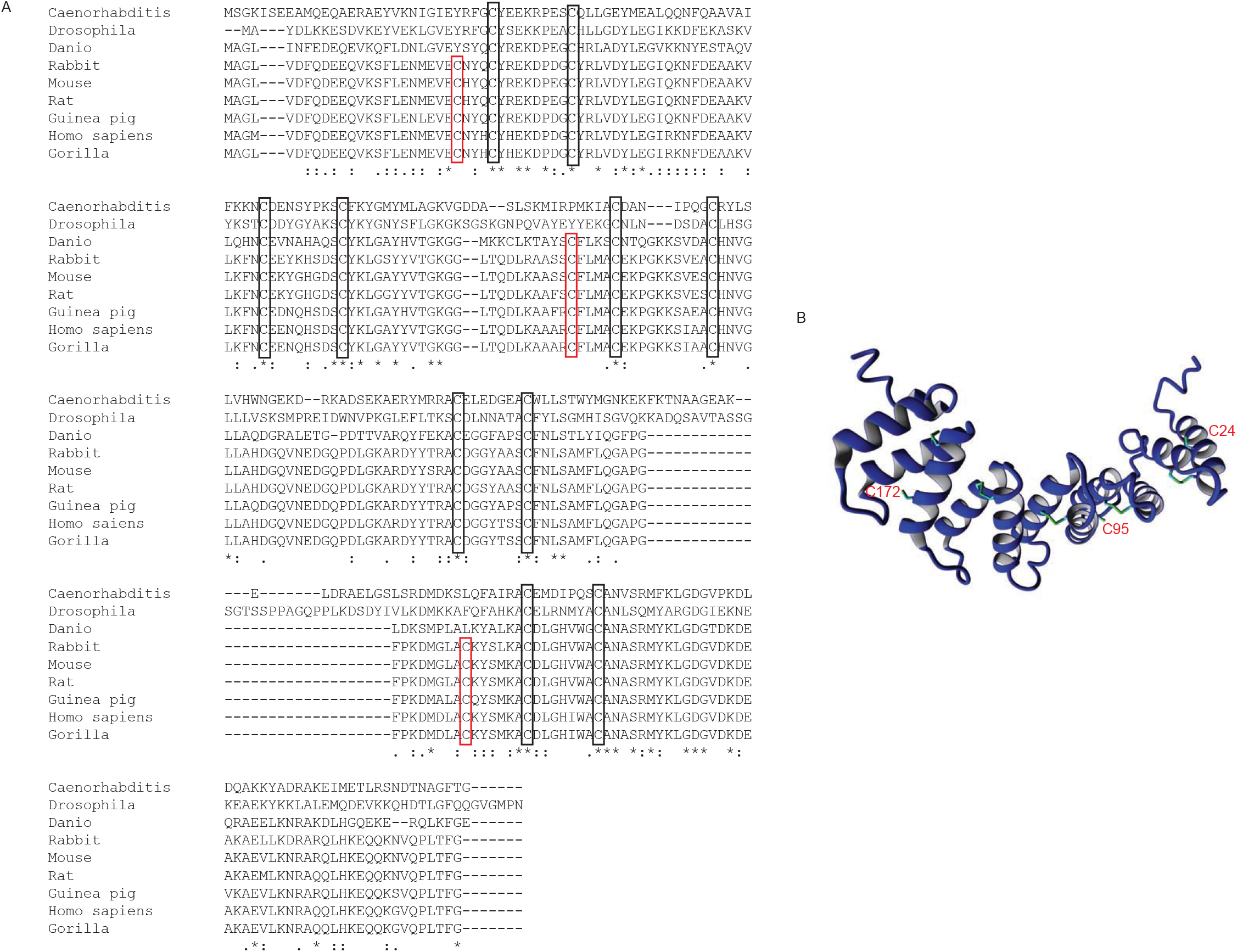
Evolutionary conservation and oxidation state of cysteine residues. (A) Multiple sequence alignment of RESA1 sequence across metazoans. Red boxes indicate cysteine residues that are present only in mammals. Black boxes indicate cysteine residues that are well conserved across eukaryotes. (B) Homology-modeled structure of RESA1 that depicts cysteine residues that are involved in disulfide bonds based on homology modeling.

**Figure EV2.**
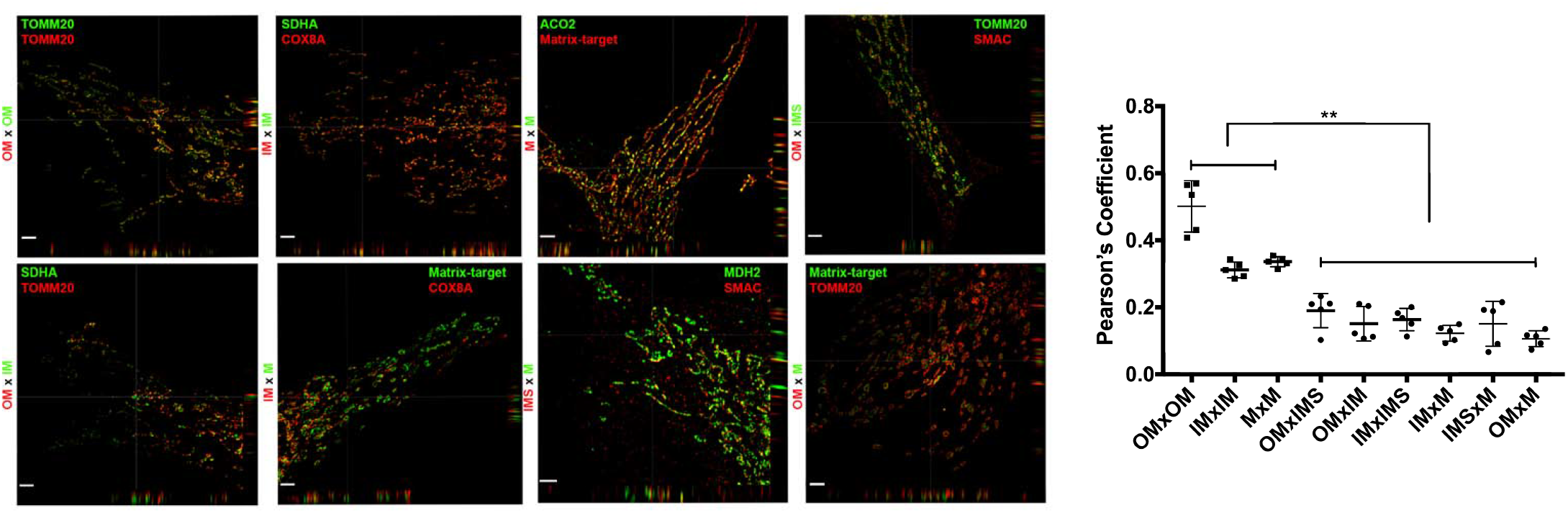
Super-resolution images for control subcompartments localization. The figure shows N-SIM super-resolution micrographs of one Z-stack (0.15 μm) orthogonal section (XYZ) of Hela cells transfected with different subcompartment markers TOMM20-DsRed (OM), COX8A-DsRed (IM), and mtPAGFP (Matrix-target) labeled with anti-TOMM20 (OM), anti-SDHA (IM), anti-Aconitase2 (M) and Smac/Diablo (IMS) antibodies. The picture represents the majority population of cells from three independent experiments. Scale bar = 2 μm. The panel shows Pearson’s coefficient in a co-localized volume of different subcompartment combinations. The data are expressed as mean ± SD (n = 5). **p < 0.01 (one-way ANOVA).

**Figure EV3.**
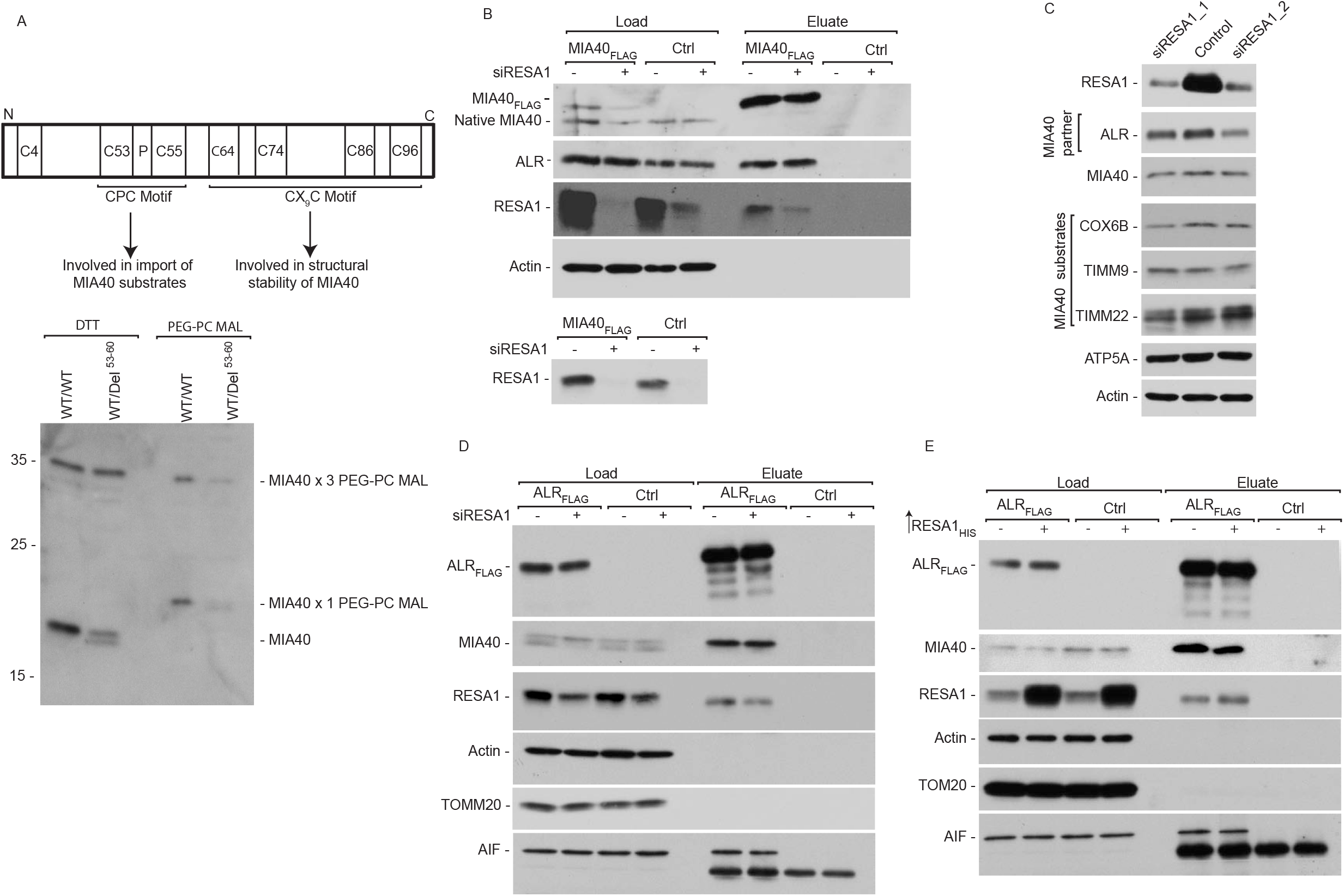
RESA1 does not affect MIA40-ALR interaction. (A) Mitochondria were isolated from HEK293 WT/WT and HEK293 MIA40 WT/Del^53-60^ cells. The mitochondria were then treated with either DTT or PEG-PC MAL, a modifying agent that binds to free cysteine residues. The reagent provides a shift of 5 kDa upon binding to each cysteine residue. The samples were analyzed by SDS-PAGE and Western blot. A schematic representation of cysteine residues and their function in MIA40 is shown. (B) Protein extracts were isolated from Flp-In 293 T-REx cells that expressed MIA40_FLAG_ that was transfected with oligonucleotides that targeted RESA1 mRNA or Mission siRNA universal negative control and subjected to affinity purification. The samples were analyzed by SDS-PAGE and Western blot. Load: 2.5%. Eluate: 100%. (C) Cellular protein extracts were isolated from HeLa cells that were transfected with oligonucleotides that targeted different regions of RESA1 mRNA or Mission siRNA universal negative control. The samples were analyzed by reducing SDS-PAGE and Western blot. (D) Protein extracts were isolated from Flp-In 293 T-REx cells that expressed ALR_FLAG_ that was transfected with oligonucleotides that targeted RESA1 mRNA or Mission siRNA universal negative control and subjected to affinity purification. The samples were analyzed by SDS-PAGE and Western blot. Load: 2.5%. Eluate: 100%. (E) Flp-In 293 T-REx cells that expressed ALR_FLAG_ were transfected with a plasmid that encoded RESA1_HIS_ or an empty vector. The affinity purification of ALR_FLAG_ was performed, and eluate fractions were analyzed by SDS-PAGE and Western blot.

**Figure EV4.**
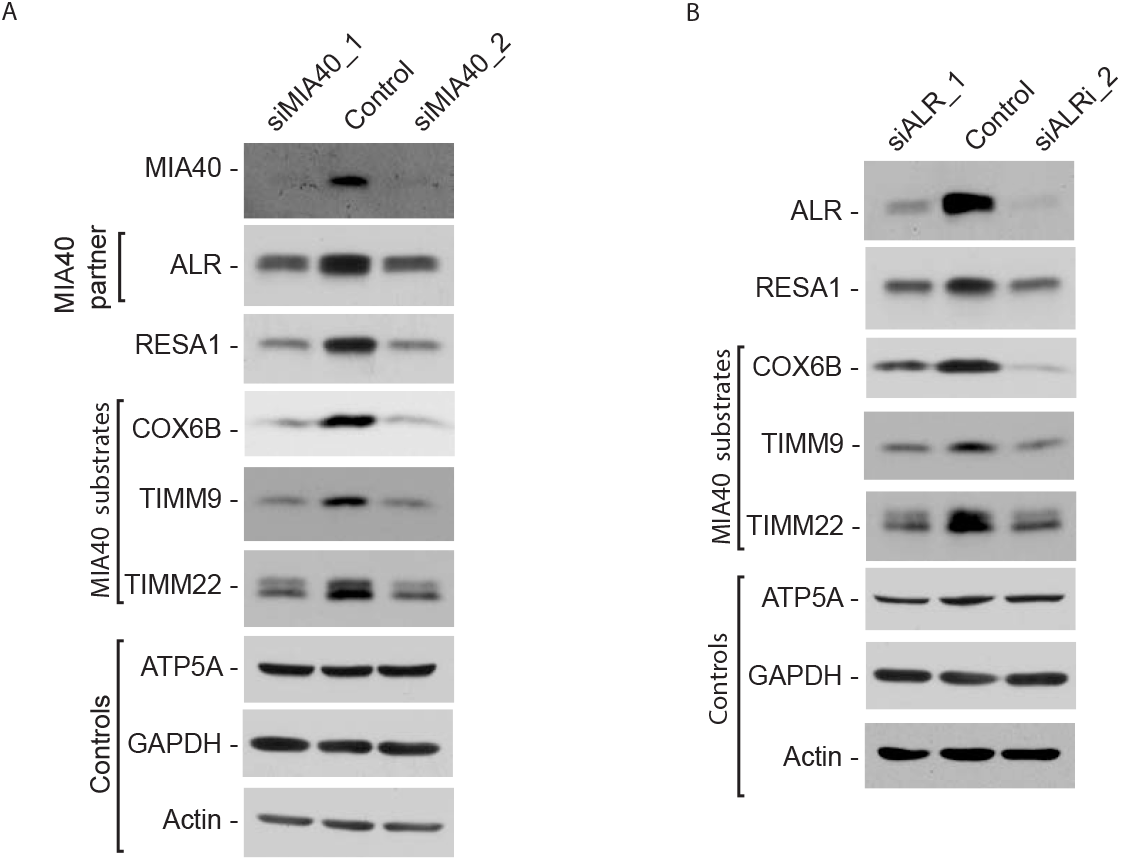
Changes in steady-state levels of RESA1 upon MIA40 or ALR silencing. (A) Cellular protein extracts were isolated from HeLa cells that were transfected with oligonucleotides that targeted different regions of MIA40 mRNA or Mission siRNA universal negative control. The samples were analyzed by reducing SDS-PAGE and Western blot. (B) Cellular protein extracts were isolated from HeLa cells that were transfected with oligonucleotides that targeted different regions of ALR mRNA or Mission siRNA universal negative control. The samples were analyzed by reducing SDS-PAGE and Western blot.

**Figure EV5.**
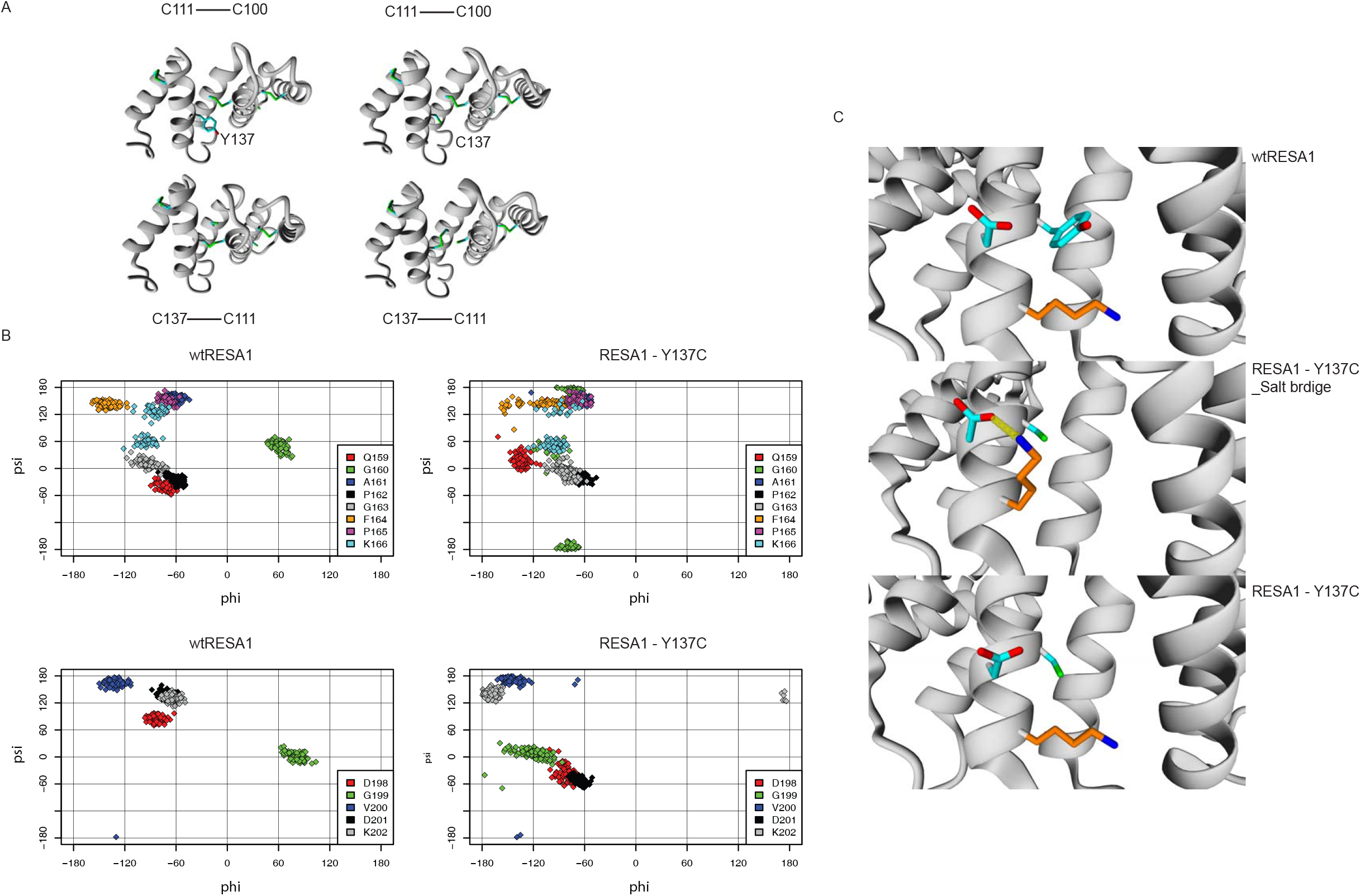
Structural modeling of wildtype and mutant RESA1. (A) Homology-modeled structure that depicts alternate disulfide bonds in the RESA1-Y137C mutant. (B) Ramachandran plot of amino acids in the loop region of the RESA1-Y137C mutant. Flexibility of the loop regions was quantified by backbone dihedral angles (phi and psi) of individual residues, variations of which were plotted in the Ramachandran plots. The conformation of G160 and G199 differed substantially upon Y137C replacement, and the conformation of Q159, D198, D201, and K202 was also altered. The analyzed residues had wider distributions of dihedral angles (phi and psi) in the RESA1-Y137C mutant, indicating greater flexibility of these loops. No significant change was observed in the conformation of the other loops in the RESA1-Y137C mutant. Wt, wildtype. (C) Salt bridge formation in the RESA1-Y137C mutant. Wildtype RESA1 and mutant RESA1 were subjected to molecular dynamics simulations, and a snapshot of salt bridge formation in the mutant is presented.

**Figure EV6.**
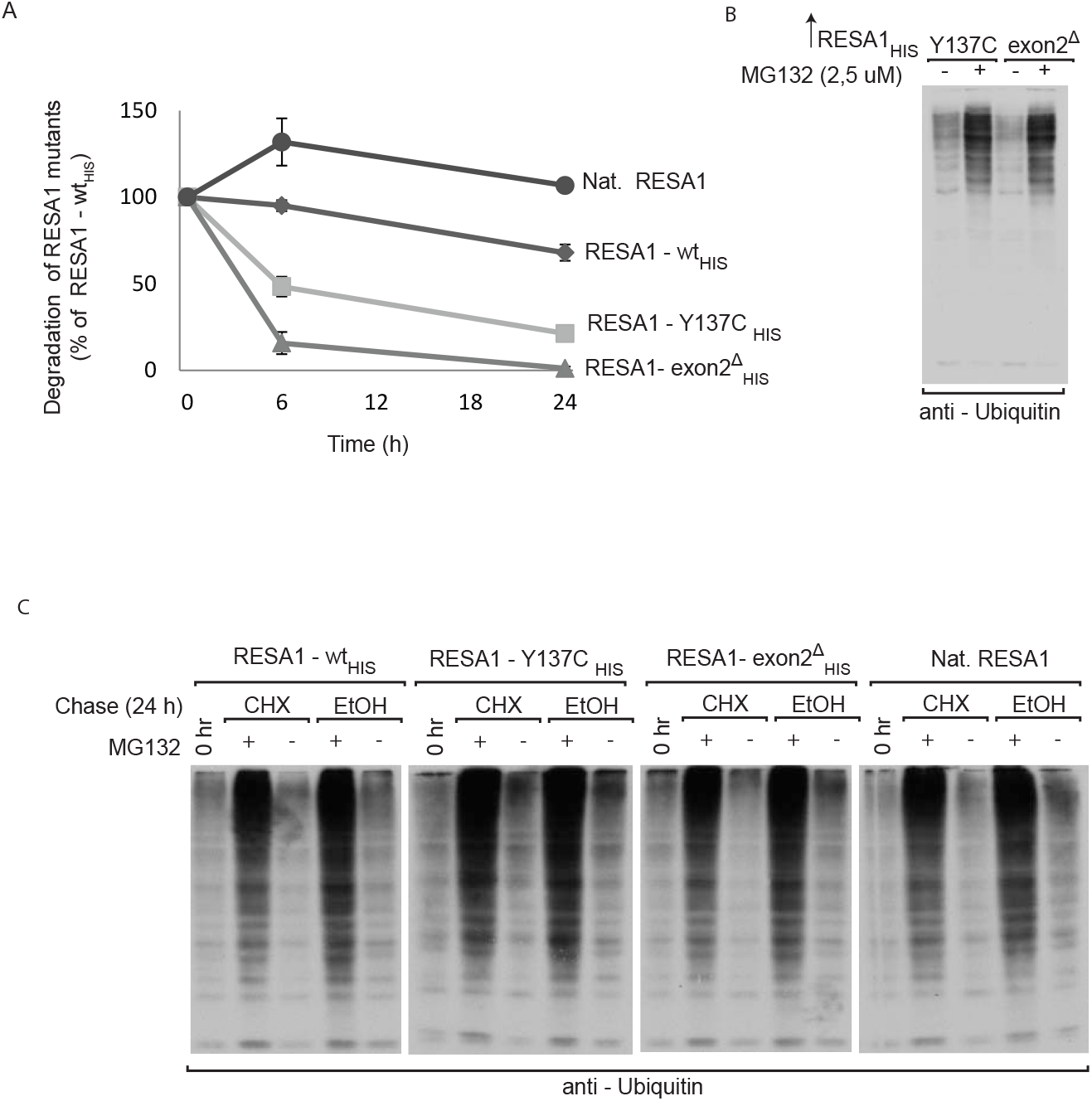
Degradation of RESA1 mutants and submitochondrial localization of RESA1-Y137C. (A) The results of three biological replicates were analyzed, quantified, and normalized to the level of native RESA1 at time 0. The data are expressed as mean ± SEM. Nat, native. (B) Samples from Fig 8E analyzed by SDS-PAGE and Western blot against ubiquitin. (C) Samples from Fig 8G analyzed by SDS-PAGE and Western blot against ubiquitin.

